# COVID-19: Variant screening, an important step towards precision epidemiology

**DOI:** 10.1101/2020.10.19.345140

**Authors:** Amrita Chattopadhyay, Tzu-Pin Lu, Ching-Yu Shih, Liang-Chuan Lai, Mong-Hsun Tsai, Eric Y. Chuang

## Abstract

Precision epidemiology using genomic technologies allows for a more targeted approach to COVID-19 control and treatment at individual and population level, and is the urgent need of the day. It enables identification of patients who may be at higher risk than others to COVID-19-related mortality, due to their genetic architecture, or who might respond better to a COVID-19 treatment. The COVID-19 virus, similar to SARS-CoV, uses the ACE2 receptor for cell entry and employs the cellular serine protease TMPRSS2 for viral S protein priming. This study aspires to present a multi-omics view of how variations in the *ACE2* and *TMPRSS2* genes affect COVID-19 infection and disease progression in affected individuals. It reports, for both genes, several variant and gene expression analysis findings, through (i) comparison analysis over single nucleotide polymorphisms (SNPs), that may account for the difference of COVID-19 manifestations among global sub-populations; (ii) calculating prevalence of structural variations (copy number variations (CNVs) / insertions), amongst populations; and (iii) studying expression patterns stratified by gender and age, over all human tissues. This work is a good first step to be followed by additional studies and functional assays towards informed treatment decisions and improved control of the infection rate.

## Introduction

The novel coronavirus of 2019 has been the cause of a global health emergency, declared as a world-wide pandemic of COVID-19, with symptoms including fever, severe respiratory illness, and pneumonia ^1^ leading to multiple organ failure and eventual sepsis ^2^, followed by death in severe cases. Studies have indicated the similarity of COVID-19 to the severe acute respiratory syndrome coronavirus (SARS-CoV) ^3,4^; however the major challenge with COVID-19 is its comparatively higher human-to-human transmission rate ^5,6^. The COVID-19 virus belongs to the beta-coronavirus genus, which also includes the highly pathogenic SARS-CoV and Middle East respiratory syndrome coronavirus (MERS-CoV) ^7^. Monumental efforts are underway to find drugs and vaccines that could potentially be used to treat people with COVID-19 as well as help prevent infection. Prior studies have revealed that the COVID-19 virus, similar to SARS-CoV, uses the angiotensin-converting enzyme 2 (ACE2) receptor for cell entry ^8^. The SARS-CoV S protein (SARS-S) engages ACE2 as the entry receptor ^9^ and employs the cellular serine protease TMPRSS2 for S protein priming ^10^. The efficiency of ACE2 was found to be a key determinant of SARS-CoV transmissibility ^11,12^. SARS-S and SARS-2-S (from COVID-19) share ~76% amino acid identity ^9^, and therefore a confirmation that SARS-2-S, like SARS-S, employs ACE2 and TMPRSS2 for host cell entry and disease progression, would provide the scientific community with relevant information regarding treatment of infected people and controlling the infection rate.

Precision epidemiology constitutes an increase in both scale and resolution of inference of genomic technologies that take a targeted approach towards infectious disease control ^13^. It includes genome-based approaches for providing information on molecular diagnosis and individual-level treatment regimens. To this end, pharmacogenetics studies involving multi-omics evidence could be a strategy to control the uncertainty of treatment decisions for severely ill COVID-19 patients. COVID-19 has a wide range of presentation, with some patients dying ^14^ while others are asymptomatic ^15^. Other than the clear risks that are associated with age and comorbidities (due to preexisting chronic conditions such as cardiac diseases, diabetes, or cancer) ^16–18^, such variability in the manifestation of symptoms and in outcomes needs to be explained through genetic probing, as drug development guided by genetic evidence should have greater rates of success ^19,20^. Identifying patients who may be at a higher risk of COVID-19-related mortality, due to their genetic architecture, or patients who might respond particularly well to a particular drug, could lead to help guide treatment decisions and successfully treat the symptoms. It could also help explain why otherwise healthy individuals in low-risk groups sometimes experience severe disease symptoms. It is necessary to identify variants of the *ACE2* and *TMPRSS2* genes that confer higher susceptibility to fatality and symptoms. However, there is the possibility that a susceptibility gene is likely to have low penetrance, and thus not all carriers will develop the disorder. Specific environmental influences are also more likely to be important risk factors that could have an effect on the severity of transmission and infections. Therefore, we will present multiple lines of supporting evidence from multi-omics data such as single nucleotide polymorphisms (SNPs), allele frequency information, structural variations such as copy number variations (CNVs i.e. deletions, duplications) and, insertions, and gene expression information for individual genes, to illustrate how multi-omics approaches can help identify COVID-19 risk factors.

Early COVID-19 data is alarming, when observed from the racial point of view. The death/recovery ratio, which is defined as the number of deaths caused by the virus divided by the number of people that were infected by it and recovered, is highly variable across ethnicities. While COVID-19 infection was observed in individuals from South and East Asian countries as early as end-of-2019 or beginning-of-2020, the recovery rates have been relatively quicker with lower morbidities, than that of Western countries such as the USA and the UK, where people appear to remain affected for longer with slower recovery times and exhibit higher death/recovery ratios ^21^. The global death toll from the coronavirus climbed to 372,116 as of June 1^st^, 2020, while the number of cases surpassed 6.17 million, according to a running tally by US-based Johns Hopkins University (https://coronavirus.jhu.edu/map.html). However, the 10 ASEAN countries (https://www.aseanbriefing.com/news/coronavirus-asia-asean-live-updates-by-country/) reported around 91,180 cases so far, with the total number of fatalities standing at 2,773. Others including Taiwan (441 confirmed cases and 7 deaths) and South Korea (11,344 confirmed cases and 269 deaths) also display very low death/recovery ratios. Densely populated developing countries in South Asia and parts of Africa have also been observed to fare far better when it comes to the mortality rate of COVID-19. The case fatality ratio (CFR) in South Asian countries such as India is 3.3%, Pakistan 2.2%, Bangladesh 1.5% and Sri Lanka 1%. Moreover, as of early April 2020, 72% of people who died of COVID-19 in Chicago, USA, were black (one-third of the city’s population), while in Georgia, as of 17 April, 40% of COVID-19 cases were white people (58% of the state) (https://dph.georgia.gov/covid-19-daily-status-report). In the UK, of the first 2,249 patients with confirmed COVID-19, 35% were non-white, which is a lot higher than the proportion of non-white people in England and Wales – 14%, according to the most recent census (http://www.ons.gov.uk/ons/dcp171778_290685.pdf). All of this data suggests a likely population level genetic variation in terms of susceptibility to the coronavirus and COVID-19 manifestations. Therefore, distribution of any genetic risk factors between populations could be the underlying mechanism leading to population-specific risk levels.

## Methods

### Single Nucleotide Polymorphism Analysis

Prior studies have suggested that genetic variants in *ACE2* might affect ACE2 levels in the human body ^22,23^. Several computational variant annotation tools are available that provide integrated reports that can be used for further rule-based filtering. VariED is one such online database developed by our research group, the first among its peers, to provide an integrated database of gene annotation and expression profiles for variants related to human diseases ^24^. VariED was utilized to obtain allele frequencies for all SNP variants from both *ACE2* and *TMPRSS2* for different global sub-populations, American (AMR), African (AFR), Finnish (FIN), Non-Finnish Europeans (NFE), South-Asian (SAS), East-Asian (EAS) and other populations (OTH). Variant and allele frequency data infers gene and variant function (e.g., whether a gene is essential) and is helpful in population genetics analyses. Although these data are collected from healthy people, such an analysis would help us screen potential targets for further functional assays or cohort studies. We therefore conducted a two-step variant allele comparison analysis over all SNPs from both genes, to obtain significant variants contributing to the difference among all populations, using Fisher’s exact test. First a general Fisher’s exact test with a Monte Carlo approximation was applied to all SNPs from both genes, over all populations, followed by a post-hoc analysis, that was conducted over the selected SNPs from the first step, to pinpoint populations that contribute the most to the variation in allele frequency through a 2X2 Fisher’s exact test. A cut-off of P < 0.05 was considered as statistically significant.

### Structural variation analysis

Knowledge of genetic structural variations in humans has accrued slowly, and studies have revealed that structural variation contributes to all classes of disease with a genetic etiology, including infectious diseases^25^. Due to the availability of CNV data in large population samples, CNVs have been the focus of studies investigating the functional consequences of structural variation. CNVs are an important and large source of both normal and pathogenic variation, and the major challenge associated with CNVs is the estimation of whether the variation is benign or affects a vital biological function and results in disease. To identify causative CNVs, the first step is to check the presence of the CNV in control cohorts and then use classifier programs or databases to predict the disease potential of the CNV. In this study, we focused on studying the CNV frequencies of our candidate genes in control cohorts and accordingly, we utilized CNVIntegrate ^26^, a web-based system developed by our research group, which is an integrated, sorted, and structured database built using CNV datasets from multiple sources, to query the genes of interest. CNV information from (i) ExAC, which consists of healthy individuals from global sub-populations, and (ii) TWCNV, a CNV database constructed from healthy Taiwanese individuals, was used to identify the prevalence of structural variations (CNVs or insertions) in genes *ACE2* and *TMPRSS2*. Thereby, we calculated the duplications and deletions frequencies for both ACE2 and TMPRSS2 among the healthy population in the TWCNV and ExAC databases. We corroborated the results from CNVIntegrate with the structural variation query results for both the genes in the GnomAD browser ^27^.

### Gene Expression Analysis

*ACE2* has been observed to be expressed predominantly by epithelial cells of the lung, intestine, kidney and blood vessels ^28^. *TMPRSS2* has been reported previously to be expressed in normal human tissues ^29^, in small intestine, prostate, colon, stomach, and salivary glands. We used a gene expression database/web tool, CellExpress ^30^, developed by our research group, to study the gene expression patterns over all tissues for both *ACE2* and *TMPRSS2*. Some studies have shown that in addition to cardiac and diabetic conditions, cancer ^31^ is associated with a high rate of comorbidity for COVID-19; 20% of the cases ending in death in Italy had a medical history of malignancy in the previous 5 years, and in Wuhan, patients (>60 years of age) with non-small cell lung cancer had the highest incidence rate for COVID-19, followed by esophageal and breast cancer ^31^. The report also showed that these patients were more likely to suffer life-threatening complications requiring emergency ICU admission or mechanical ventilation, with death attributed to acute myocardial infarction, acute respiratory syndrome, septic shock, and pulmonary embolism. Therefore, we conducted a case control analysis for both *ACE2* and *TMPRSS2* using CellExpress, with the case data for 36 cancer types acquired from GSE36133 (Cancer Cell line Encyclopedia) ^32^, and the control data from the Roth normal dataset (GSE7307). We also did a gene expression search for both genes over all tissues in cancer datasets, stratified by gender and age, using CellExpress.

## Results

### SNP analysis results

After filtering out the uninformative SNPs with no reported allele frequencies, a total of 362 SNPs from *ACE2* and 532 SNPs from *TMPRSS2* were used to compare the allele frequencies among different sub-populations from the Exome Aggregation Consortium (ExAC) Database ^33^. In the first step, 44 significant SNPs from *ACE2* and 98 SNPs from *TMPRSS2* were obtained from the Fisher’s exact test with a Monte Carlo approximation, that were found to display different allele frequencies between sub-populations. Then a 2X2 Fisher’s exact test on selected SNPs from the first step to identify populations that contribute the most to the variation in allele frequency. Table 1 and Table 2 display selected SNPs with significant allele frequency variation. The reported SNPs were found to have higher allele frequencies among Africans and East Asian populations compared to Europeans and Americans. Africans were found to display particularly significant variation for most of the reported SNPs from both *ACE2* and *TMPRSS2*, which may suggest differential susceptibility towards coronavirus in the respective populations under similar conditions. Most of the reported SNPs in this study have also been reported in other COVID-19-related studies ^34–39^. Supplementary file 1 and Supplementary file 2 give a full list of all the significant findings.

**Table 1.**
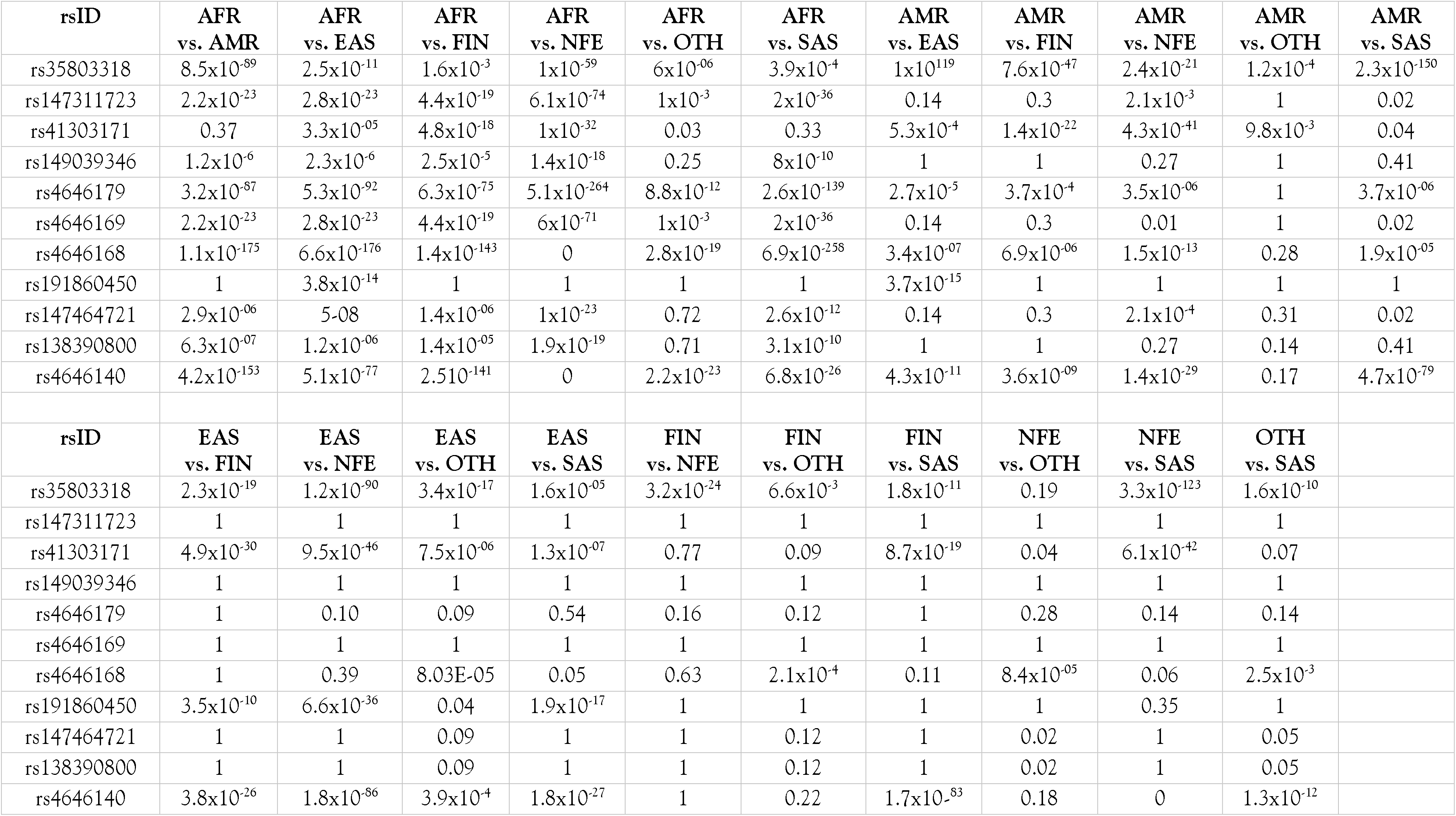
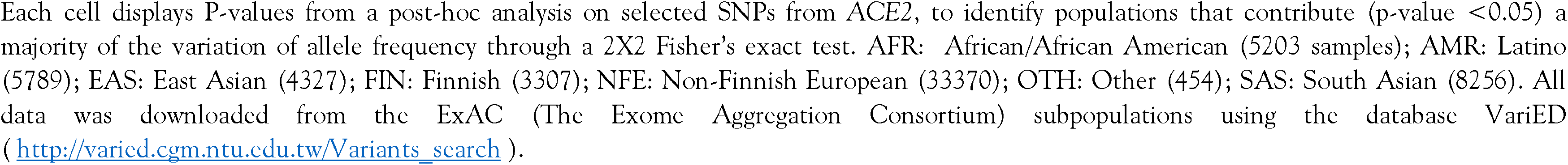
SNPs from *ACE2* with significant allele frequency variation between ethnic populations. Description: Each cell displays P-values from a post-hoc analysis on selected SNPs from *ACE2*, to identify populations that contribute (p-value <0.05) a majority of the variation of allele frequency through a 2X2 Fisher’s exact test. AFR: African/African American (5203 samples); AMR: Latino (5789); EAS: East Asian (4327); FIN: Finnish (3307); NFE: Non-Finnish European (33370); OTH: Other (454); SAS: South Asian (8256). All data was downloaded from the ExAC (The Exome Aggregation Consortium) subpopulations using the database VariED (http://varied.cgm.ntu.edu.tw/Variants_search).

| rsID | AFR<br>vs. AMR | AFR<br>vs. EAS | AFR<br>vs. FIN | AFR<br>vs. NFE | AFR<br>vs. OTH | AFR<br>vs. SAS | AMR<br>vs. EAS | AMR<br>vs. FIN | AMR<br>vs. NFE | AMR<br>vs. OTH | AMR<br>vs. SAS |
| --- | --- | --- | --- | --- | --- | --- | --- | --- | --- | --- | --- |
| rs35803318 | 8.5x10<sup>-89</sup> | 2.5x10<sup>-11</sup> | 1.6x10<sup>-3</sup> | 1x10<sup>-59</sup> | 6x10<sup>-06</sup> | 3.9x10<sup>-4</sup> | 1x10<sup>-119</sup> | 7.6x10<sup>-47</sup> | 2.4x10<sup>-21</sup> | 1.2x10<sup>-4</sup> | 2.3x10<sup>-150</sup> |
| rs147311723 | 2.2x10<sup>-23</sup> | 2.8x10<sup>-23</sup> | 4.4x10<sup>-19</sup> | 6.1x10<sup>-74</sup> | 1x10<sup>-3</sup> | 2x10<sup>-36</sup> | 0.14 | 0.3 | 2.1x10<sup>-3</sup> | 1 | 0.02 |
| rs41303171 | 0.37 | 3.3x10<sup>-05</sup> | 4.8x10<sup>-18</sup> | 1x10<sup>-32</sup> | 0.03 | 0.33 | 5.3x10<sup>-4</sup> | 1.4x10<sup>-22</sup> | 4.3x10<sup>-41</sup> | 9.8x10<sup>-3</sup> | 0.04 |
| rs149039346 | 1.2x10<sup>-6</sup> | 2.3x10<sup>-6</sup> | 2.5x10<sup>-5</sup> | 1.4x10<sup>-18</sup> | 0.25 | 8x10<sup>-10</sup> | 1 | 1 | 0.27 | 1 | 0.41 |
| rs4646179 | 3.2x10<sup>-87</sup> | 5.3x10<sup>-92</sup> | 6.3x10<sup>-75</sup> | 5.1x10<sup>-264</sup> | 8.8x10<sup>-12</sup> | 2.6x10<sup>-139</sup> | 2.7x10<sup>-5</sup> | 3.7x10<sup>-4</sup> | 3.5x10<sup>-06</sup> | 1 | 3.7x10<sup>-06</sup> |
| rs4646169 | 2.2x10<sup>-23</sup> | 2.8x10<sup>-23</sup> | 4.4x10<sup>-19</sup> | 6x10<sup>-71</sup> | 1x10<sup>-3</sup> | 2x10<sup>-36</sup> | 0.14 | 0.3 | 0.01 | 1 | 0.02 |
| rs4646168 | 1.1x10<sup>-175</sup> | 6.6x10<sup>-176</sup> | 1.4x10<sup>-143</sup> | 0 | 2.8x10<sup>-19</sup> | 6.9x10<sup>-258</sup> | 3.4x10<sup>-07</sup> | 6.9x10<sup>-06</sup> | 1.5x10<sup>-13</sup> | 0.28 | 1.9x10<sup>-05</sup> |
| rs191860450 | 1 | 3.8x10<sup>-14</sup> | 1 | 1 | 1 | 1 | 3.7x10<sup>-15</sup> | 1 | 1 | 1 | 1 |
| rs147464721 | 2.9x10<sup>-06</sup> | 5-08 | 1.4x10<sup>-06</sup> | 1x10<sup>-23</sup> | 0.72 | 2.6x10<sup>-12</sup> | 0.14 | 0.3 | 2.1x10<sup>-4</sup> | 0.31 | 0.02 |
| rs138390800 | 6.3x10<sup>-07</sup> | 1.2x10<sup>-06</sup> | 1.4x10<sup>-05</sup> | 1.9x10<sup>-19</sup> | 0.71 | 3.1x10<sup>-10</sup> | 1 | 1 | 0.27 | 0.14 | 0.41 |
| rs4646140 | 4.2x10<sup>-153</sup> | 5.1x10<sup>-77</sup> | 2.51x10<sup>-141</sup> | 0 | 2.2x10<sup>-23</sup> | 6.8x10<sup>-26</sup> | 4.3x10<sup>-11</sup> | 3.6x10<sup>-09</sup> | 1.4x10<sup>-29</sup> | 0.17 | 4.7x10<sup>-79</sup> |
| rsID | EAS<br>vs. FIN | EAS<br>vs. NFE | EAS<br>vs. OTH | EAS<br>vs. SAS | FIN<br>vs. NFE | FIN<br>vs. OTH | FIN<br>vs. SAS | NFE<br>vs. OTH | NFE<br>vs. SAS | OTH<br>vs. SAS |  |
| rs35803318 | 2.3x10<sup>-19</sup> | 1.2x10<sup>-90</sup> | 3.4x10<sup>-17</sup> | 1.6x10<sup>-05</sup> | 3.2x10<sup>-24</sup> | 6.6x10<sup>-3</sup> | 1.8x10<sup>-11</sup> | 0.19 | 3.3x10<sup>-123</sup> | 1.6x10<sup>-10</sup> |  |
| rs147311723 | 1 | 1 | 1 | 1 | 1 | 1 | 1 | 1 | 1 | 1 |  |
| rs41303171 | 4.9x10<sup>-30</sup> | 9.5x10<sup>-46</sup> | 7.5x10<sup>-06</sup> | 1.3x10<sup>-07</sup> | 0.77 | 0.09 | 8.7x10<sup>-19</sup> | 0.04 | 6.1x10<sup>-42</sup> | 0.07 |  |
| rs149039346 | 1 | 1 | 1 | 1 | 1 | 1 | 1 | 1 | 1 | 1 |  |
| rs4646179 | 1 | 0.10 | 0.09 | 0.54 | 0.16 | 0.12 | 1 | 0.28 | 0.14 | 0.14 |  |
| rs4646169 | 1 | 1 | 1 | 1 | 1 | 1 | 1 | 1 | 1 | 1 |  |
| rs4646168 | 1 | 0.39 | 8.03E-05 | 0.05 | 0.63 | 2.1x10<sup>-4</sup> | 0.11 | 8.4x10<sup>-05</sup> | 0.06 | 2.5x10<sup>-3</sup> |  |
| rs191860450 | 3.5x10<sup>-10</sup> | 6.6x10<sup>-36</sup> | 0.04 | 1.9x10<sup>-17</sup> | 1 | 1 | 1 | 1 | 0.35 | 1 |  |
| rs147464721 | 1 | 1 | 0.09 | 1 | 1 | 0.12 | 1 | 0.02 | 1 | 0.05 |  |
| rs138390800 | 1 | 1 | 0.09 | 1 | 1 | 0.12 | 1 | 0.02 | 1 | 0.05 |  |
| rs4646140 | 3.8x10<sup>-26</sup> | 1.8x10<sup>-86</sup> | 3.9x10<sup>-4</sup> | 1.8x10<sup>-27</sup> | 1 | 0.22 | 1.7x10<sup>-83</sup> | 0.18 | 0 | 1.3x10<sup>-12</sup> |  |

**Table 2.** SNPs from *TMPRSS2* with significant allele frequency variation between ethnic populations. Description: Each cell displays P-values from a post-hoc analysis on selected SNPs from *TMPRSS2*, to identify populations that contribute (p-value <0.05) a majority of the variation of allele frequency through a 2X2 Fisher’s exact test. AFR: African/African American (5203 samples); AMR: Latino (5789); EAS: East Asian (4327); FIN: Finnish (3307); NFE: Non-Finnish European (33370); OTH: Other (454); SAS: South Asian (8256). All data was downloaded from the ExAC (The Exome Aggregation Consortium) subpopulations using the database VariED (http://varied.cgm.ntu.edu.tw/Variants_search).

| rs_ID | AFR<br>vs. AMR | AFR<br>vs. EAS | AFR<br>vs. FIN | AFR<br>vs. NFE | AFR<br>vs. OTH | AFR<br>vs. SAS | AMR<br>vs. EAS | AMR<br>vs. FIN | AMR<br>vs. NFE | AMR<br>vs. OTH | AMR<br>vs. SAS |
| --- | --- | --- | --- | --- | --- | --- | --- | --- | --- | --- | --- |
| rs12329760 | $1 \times 10^{-65}$ | $6 \times 10^{-21}$ | $1 \times 10^{-15}$ | $5 \times 10^{-20}$ | 0.51 | $3 \times 10^{-09}$ | $1 \times 10^{-147}$ | $2 \times 10^{-120}$ | $4 \times 10^{-40}$ | $6 \times 10^{-10}$ | $2 \times 10^{-38}$ |
| rs3787950 | $4 \times 10^{-146}$ | $6 \times 10^{-09}$ | $1 \times 10^{-128}$ | $2 \times 10^{-134}$ | $2 \times 10^{-06}$ | 0.6 | $3 \times 10^{-79}$ | $2 \times 10^{-3}$ | $1 \times 10^{-23}$ | $1 \times 10^{-08}$ | $2 \times 10^{-167}$ |
| rs118028230 | 0.37 | $9 \times 10^{-136}$ | 1 | 0.35 | 0.001 | 0.06 | $3 \times 10^{-140}$ | 0.30 | $5 \times 10^{-3}$ | 0.01 | 0.42 |
| rs61735795 | $1 \times 10^{-05}$ | $1 \times 10^{-4}$ | $8 \times 10^{-4}$ | $1 \times 10^{-12}$ | 0.62 | $6 \times 10^{-07}$ | 1 | 1 | 1 | 1 | 1 |
| rs777860329 | $9 \times 10^{-09}$ | $1 \times 10^{-05}$ | $1 \times 10^{-10}$ | $8 \times 10^{-41}$ | 1 | $9 \times 10^{-20}$ | 0.34 | 0.05 | $2 \times 10^{-07}$ | 1 | $8 \times 10^{-4}$ |
| rs75168613 | $3 \times 10^{-159}$ | $1 \times 10^{-181}$ | $1 \times 10^{-153}$ | 0.22 | $1 \times 10^{-12}$ | $1 \times 10^{-213}$ | $3 \times 10^{-11}$ | $7 \times 10^{-12}$ | $1 \times 10^{-08}$ | $5 \times 10^{-4}$ | 0.11 |
| rs113288437 | $8 \times 10^{-174}$ | $4 \times 10^{-233}$ | $4 \times 10^{-196}$ | 0.32 | $1 \times 10^{-17}$ | $2 \times 10^{-275}$ | $1 \times 10^{-23}$ | $8 \times 10^{-23}$ | $1 \times 10^{-27}$ | 0.05 | $2 \times 10^{-07}$ |
| rs140141551 | $7 \times 10^{-12}$ | $6 \times 10^{-05}$ | $9 \times 10^{-06}$ | $2 \times 10^{-54}$ | $2 \times 10^{-08}$ | 0.11 | $1 \times 10^{-22}$ | 0.11 | $1 \times 10^{-18}$ | 0.02 | $7 \times 10^{-10}$ |
| rs2298658 | $4 \times 10^{-07}$ | $2 \times 10^{-07}$ | 0.38 | 1 | 1 | 1 | 0.75 | $4 \times 10^{-4}$ | $7 \times 10^{-16}$ | 0.4 | $1 \times 10^{-08}$ |
| rs2298659 | $2 \times 10^{-24}$ | $7 \times 10^{-18}$ | $4 \times 10^{-84}$ | $4 \times 10^{-08}$ | $1 \times 10^{-3}$ | 0.30 | 0.37 | $2 \times 10^{-27}$ | $1 \times 10^{-15}$ | 0.42 | $4 \times 10^{-25}$ |
| rs17854725 | 0.83 | $5 \times 10^{-178}$ | $6 \times 10^{-36}$ | $3 \times 10^{-91}$ | $2 \times 10^{-05}$ | $5 \times 10^{-19}$ | $6 \times 10^{-183}$ | $2 \times 10^{-38}$ | $1 \times 10^{-102}$ | $1 \times 10^{-05}$ | $3 \times 10^{-21}$ |
| rs74423429 | 0.88 | $6 \times 10^{-06}$ | $2 \times 10^{-25}$ | $2 \times 10^{-31}$ | $3 \times 10^{-3}$ | 0.01 | $1 \times 10^{-05}$ | $1 \times 10^{-27}$ | $1 \times 10^{-35}$ | $2 \times 10^{-3}$ | $5 \times 10^{-3}$ |
| rs765703243 | $1 \times 10^{-11}$ | $6 \times 10^{-09}$ | $5 \times 10^{-05}$ | $1 \times 10^{-05}$ | 0.60 | $1 \times 10^{-18}$ | 0.73 | $2 \times 10^{-24}$ | $6 \times 10^{-06}$ | $1 \times 10^{-4}$ | 0.16 |
| rs_ID | EAS<br>vs. FIN | EAS<br>vs. NFE | EAS<br>vs. OTH | EAS<br>vs. SAS | FIN<br>vs. NFE | FIN<br>vs. OTH | FIN<br>vs. SAS | NFE<br>vs. OTH | NFE<br>vs. SAS | OTH<br>vs. SAS |  |
| rs12329760 | 0.53 | $4 \times 10^{-95}$ | $6 \times 10^{-06}$ | $5 \times 10^{-57}$ | $8 \times 10^{-69}$ | $3 \times 10^{-5}$ | $1 \times 10^{-43}$ | 0.03 | 0.01 | 0.14 | |
| rs3787950 | $2 \times 10^{-77}$ | $5 \times 10^{-50}$ | 0.02 | $4 \times 10^{-09}$ | $1 \times 10^{-28}$ | $2 \times 10^{-12}$ | $3 \times 10^{-139}$ | 0.01 | $2 \times 10^{-182}$ | $5 \times 10^{-06}$ | |
| rs118028230 | $1 \times 10^{-99}$ | 0.58 | $5 \times 10^{-14}$ | $4 \times 10^{-168}$ | 1 | $1 \times 10^{-3}$ | 0.07 | $2 \times 10^{-05}$ | $4 \times 10^{-06}$ | 0.02 | |
| rs61735795 | 1 | 1 | 1 | 1 | 1 | 1 | 1 | 1 | 1 | 1 |  |
| rs777860329 | $6 \times 10^{-3}$ | $3 \times 10^{-10}$ | 1 | $2 \times 10^{-5}$ | 1 | 1 | 1 | 1 | 1 | 1 | |
| rs75168613 | 0.26 | $1 \times 10^{-4}$ | $2 \times 10^{-12}$ | $1 \times 10^{-08}$ | $8 \times 10^{-06}$ | $1 \times 10^{-13}$ | $1 \times 10^{-09}$ | $8 \times 10^{-09}$ | $3 \times 10^{-05}$ | $3 \times 10^{-05}$ | |
| rs113288437 | 0.26 | $2 \times 10^{-05}$ | $2 \times 10^{-13}$ | $7 \times 10^{-11}$ | $1 \times 10^{-06}$ | $1 \times 10^{-14}$ | $1 \times 10^{-11}$ | $8 \times 10^{-09}$ | $2 \times 10^{-06}$ | $5 \times 10^{-05}$ | |
| rs140141551 | $1 \times 10^{-14}$ | $1 \times 10^{-65}$ | $4 \times 10^{-15}$ | $1 \times 10^{-08}$ | $3 \times 10^{-17}$ | $3 \times 10^{-3}$ | $4 \times 10^{-4}$ | 0.7 | $1 \times 10^{-67}$ | $8 \times 10^{-07}$ | |
| rs2298658 | $3 \times 10^{-4}$ | $7 \times 10^{-15}$ | 0.24 | $2 \times 10^{-08}$ | 0.37 | 1 | 0.49 | 1 | 1 | 1 | |
| rs2298659 | $4 \times 10^{-28}$ | $3 \times 10^{-09}$ | 0.66 | $2 \times 10^{-17}$ | $2 \times 10^{-84}$ | $5 \times 10^{-08}$ | $6 \times 10^{-91}$ | 0.12 | $2 \times 10^{-07}$ | $4 \times 10^{-3}$ | |
| rs17854725 | $1 \times 10^{-304}$ | 0.54 | $5 \times 10^{-64}$ | 0 | 0.19 | 0.15 | $4 \times 10^{-09}$ | 0.04 | $5 \times 10^{-32}$ | 0.31 | |
| rs74423429 | $1 \times 10^{-39}$ | $5 \times 10^{-47}$ | $5 \times 10^{-08}$ | $2 \times 10^{-11}$ | 0.03 | 0.08 | $4 \times 10^{-21}$ | 0.3 | $3 \times 10^{-30}$ | 0.05 | |
| rs765703243 | $1 \times 10^{-19}$ | $4 \times 10^{-4}$ | $6 \times 10^{-4}$ | 0.05 | $1 \times 10^{-20}$ | 0.26 | $1 \times 10^{-34}$ | 0.04 | $3 \times 10^{-12}$ | $2 \times 10^{-06}$ | |

### Structural Variation analysis results

The duplication and deletion frequencies for the *ACE2* and *TMPRSS2* genes were obtained from the control populations in Taiwan (TWCNV) and other global ethnic populations (ExAC) using CNVIntegrate. An intuitive model suggests that an increase in the copy number of a specific gene will, on average, lead to corresponding increase in the expression level of that gene, and vice versa ^40^. The results were consistent with this hypothesis. No duplications or deletions were observed for *ACE2* among the healthy population in the TWCNV and ExAC databases (Figure 1A). For *TMPRSS2*, duplication was observed in 0.06% and 0.014% of the samples from TWCNV and ExAC, respectively, while deletion was observed at 1.08% in TWCNV and 0.0% in ExAC (Figure 1B). The corroborated results from GnomAD browser with the structural variation query results for both the genes are displayed in Table 3. *ACE2* displayed only one duplication event, and *TMPRSS2* displayed 4 duplication/insertion and deletion events (Table 3). This suggests that these variations are rare among healthy cohorts and provide an evidence of the possibility that they could be potentially pathogenic. Moreover, whether the CNV is of clinical consequence may also depend on other factors, such as ethnic background (with specific genetic makeup), environmental elements (such as social distancing measures), age, or sex ^41^. Perception of the clinical consequences can change over time, as our knowledge grows. Moreover, the consideration of x-linked CNVs (ACE2 is located at chrX:15579156-15620271) in males is important, as many of the reported benign variants included in databases are seen in females. However, in men, who have only one X chromosome, the same change may be fundamentally pathogenic. This might also be an explanation of why more deaths from COVID-19 are observed in males than females ^42^.

**Figure 1.**
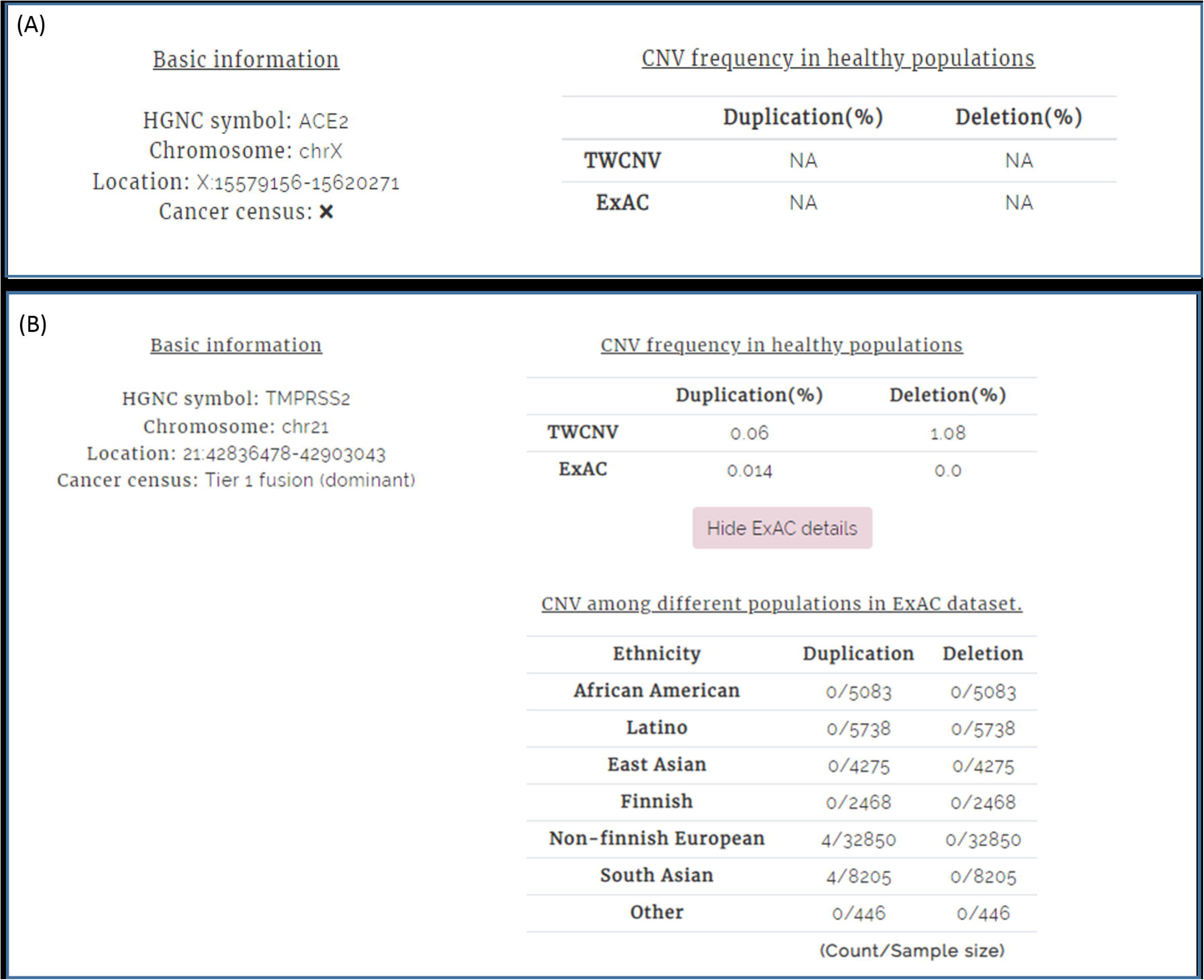
Copy number variation (CNV) query results for genes *ACE2* and *TMPRSS2* using healthy populations from the CNVIntegrate database (http://cnvintegrate.cgm.ntu.edu.tw/). (A) CNV frequency in healthy populations TWCNV and ExAC for gene *ACE2*. (B) CNV frequency in healthy populations TWCNV and ExAC (sub-populations) for gene *TMPRSS2*.

**Table 3.** Structural variation observed for *ACE2* and *TMPRSS2* in the gnomAD database.

| Gene | Variant ID | Consequence | Class | Position | Size | Allele Count | Allele Number | Allele Frequency | Homozygote Count |
| --- | --- | --- | --- | --- | --- | --- | --- | --- | --- |
| ACE2 | DUP_X_52669 | copy gain | duplication | 15377083-15726738 | 349655 | 4 | 15814 | 0.00025294 | 0 |
| TMPRSS2 | DUP_21_50235 | partial duplication | duplication | 42846101-42846693 | 592 | 36 | 21434 | 0.00168 | 0 |
| TMPRSS2 | DEL_21_180588 | intronic | deletion | 42847455-42847523 | 68 | 2969 | 21558 | 0.137721 | 0 |
| TMPRSS2 | INS_21_113715 | intronic | insertion | 42865560 | 279 | 1 | 21694 | 0.000046 | 0 |
| TMPRSS2 | DEL_21_180589 | intronic | deletion | 42869398-42869694 | 296 | 1 | 21694 | 0.000046 | 0 |

### Gene Expression Analysis results

Figure 2 displays the gene expression of baseline healthy populations from CellExpress, stratified by gender, to observe its effect (if any) on the baseline gene expression levels. Other than the pituitary gland, the baseline data did not show any significant difference in gene expression levels between males and females. Similarly, no effect was observed when gene expression was stratified by age (Figure 3). The overall results resonated with prior findings, as both the genes are found to be expressed consistently in all tissues, which was further validated by tissue-specific gene expression in the GTEx population ^43^ (Supplementary file 3). As comorbidity for cancer patients were higher, a case control analysis for both *ACE2* and *TMPRSS2* using CellExpress was conducted which demonstrated *ACE2* expression to be significantly associated with cancer, with a p-value of 7.66×10^−4^, while *TMPRSS2* expression displayed a p-value of 7.83×10^−2^. The results for a gene expression search for both genes over all tissues in cancer datasets, stratified by gender and age, using CellExpress, displayed high gene expression in all tissues for both genes, with no effect of gender and age (Supplementary file 4, Supplementary file 5).

**Figure 2.**
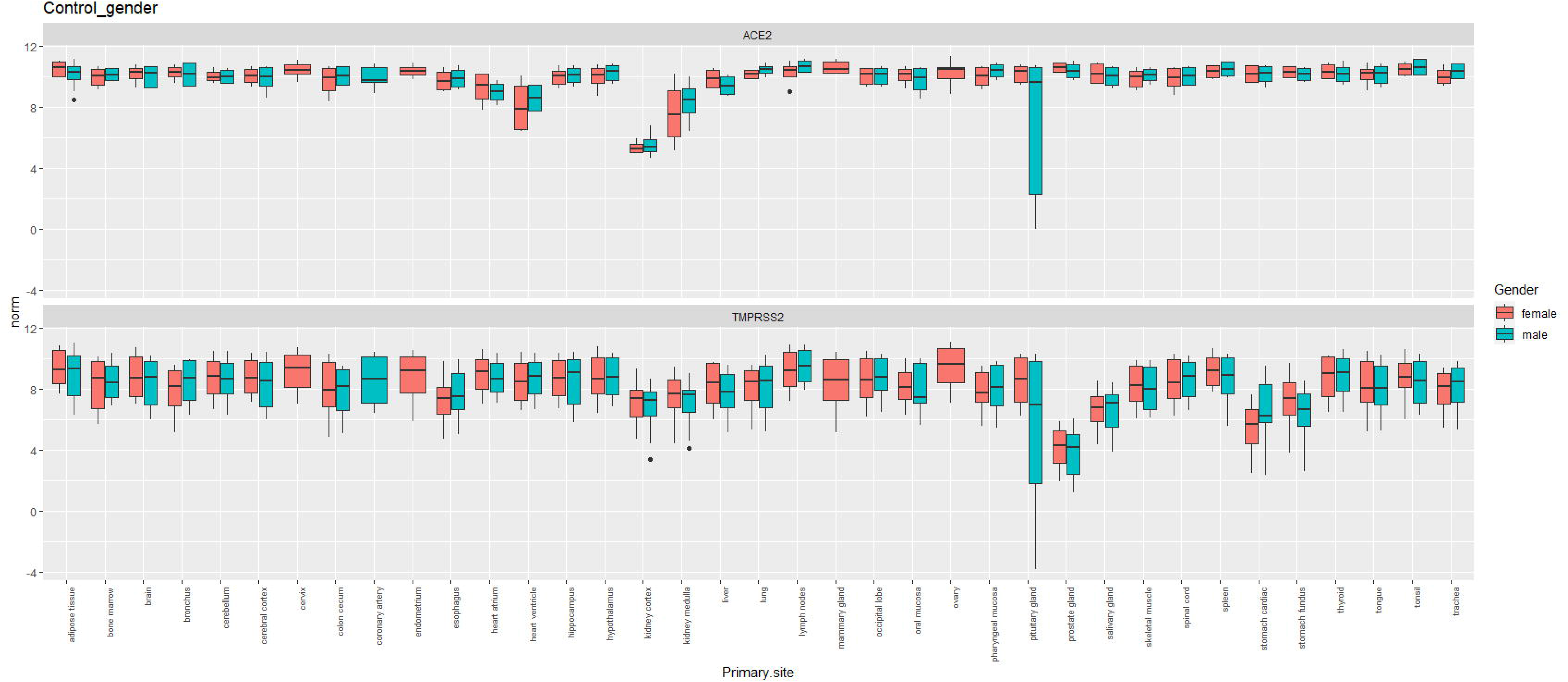
Gene expression of baseline healthy populations across different tissues from CellExpress, stratified by gender. The x-axis displays tissue names; the y-axis displays norm = log_2_ normalized gene expression values. Pink box-plots display gene expression for healthy females; blue box-plots display gene expression for healthy males. The upper panel shows gene expression for *ACE2*; the lower panel shows gene expression for *TMPRSS2*. All gene expression data were downloaded from CellExpress (http://cellexpress.cgm.ntu.edu.tw/).

**Figure 3.**
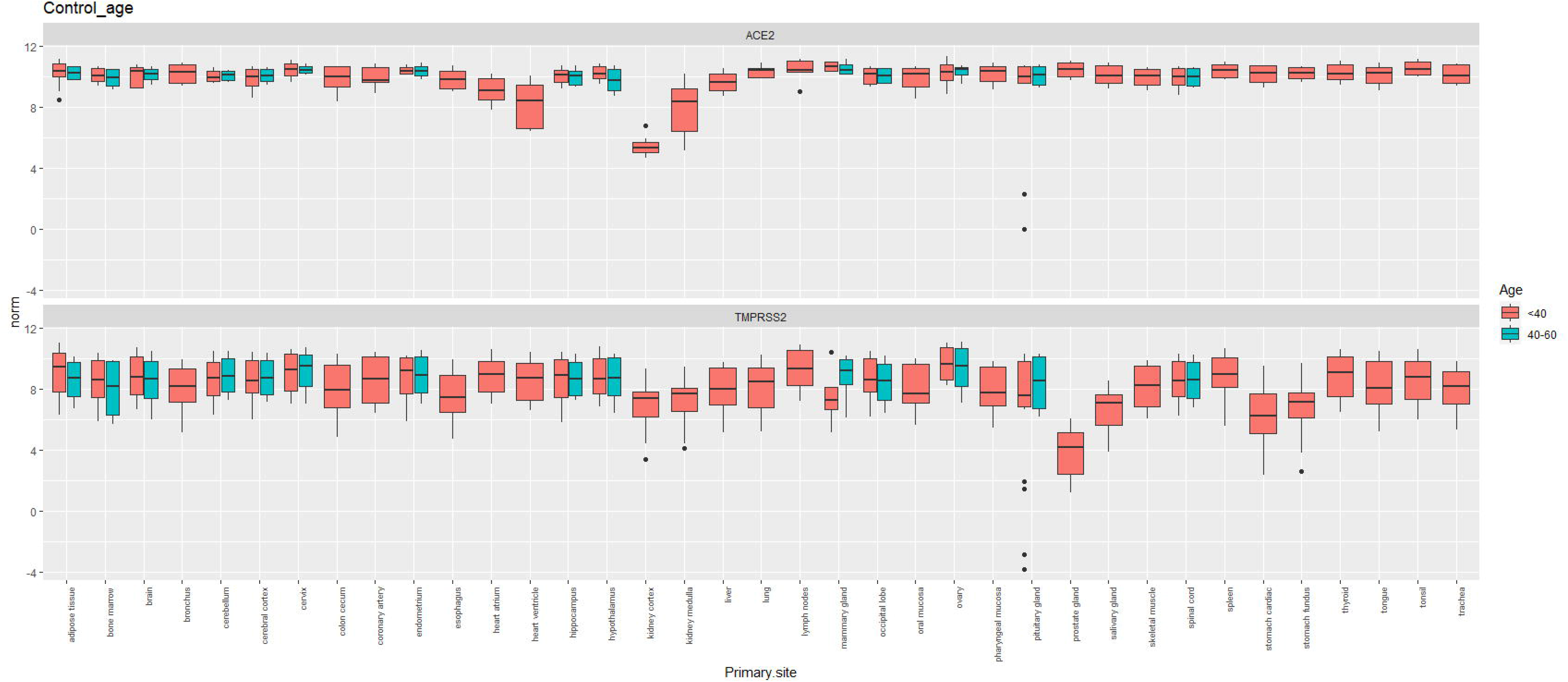
Gene expression of baseline healthy populations across different tissues from CellExpress, stratified by age. The x-axis displays tissue names; the y-axis displays norm = log_2_ normalized gene expression values. Pink box-plots display gene expression for healthy individuals <40 years of age; blue box-plots display gene expression for healthy individuals 40 - 60 years of age. The upper panel shows gene expression for *ACE2*; the lower panel shows gene expression for *TMPRSS2*. All gene expression data were downloaded from CellExpress (http://cellexpress.cgm.ntu.edu.tw/).

## Discussion

Incorporating informative priors based on biological knowledge or predicted variant function, along with integrated gene expression or other omics data, may help to inform treatment decisions for coronavirus-infected people showing COVID-19 symptoms and control the infection rate. This work provides a basis for future investigations of *ACE2* and *TMPRSS2* through further functional assays and protein expression analysis. More information on variants (SNPS and CNVs) needs to be accumulated through (1) fine mapping analysis of the variants that have been obtained through initial analysis in this study, in order to confirm their contribution to the differential responses to COVID-19 and mortality across different ethnicities in COVID-19 patients; (2) improved risk prediction accuracy through large case-control studies involving candidate gene studies and gene-gene interaction studies using genome-wide data; and (3) experimental validation of all significant findings. Gene expression analysis showed lung tissues to have comparatively higher expression of *ACE2* and *TMPRSS2*, consistent with the fact that COVID-19 affects lungs and lung tissues. Further protein expression studies are required to confirm findings in lung and different tissues. It is necessary to validate mRNA findings at the protein level, as mRNA expression patterns of *ACE2* and *TMPRSS2* are not necessarily the same as the protein expression patterns due to some post-translational modification. Furthermore, a comparison of *ACE2* and *TMPRSS2* expression levels between patients of different ethnicities also needs to be expanded to account for the various levels of affected cases and morbidity. Immune responses in infected patients play an important role in fighting coronavirus, even though immunopathological damage can occur via the cytokine storm. Extensive experimental analysis is necessary to tease out further correlations between expression of the ACE2 receptor and immune signatures in the lungs. One conjecture has been about levels of BCG (Bacille Calmette Guérin) vaccination for tuberculosis, where a striking negative correlation has been observed with COVID-19 casualties. The primary function of the BCG vaccine involves boosting immunity, and it might have a role to play in the greater immunity against the SARS-CoV-2 virus. According to the BCG world atlas, the UK and western Europe had BCG vaccination policies in the past, while in the US it is not mandatory for all groups of people. However, in Asia and eastern Europe, BCG vaccination is mandatory for all groups of people. In spite of Spain and Portugal sharing a border, the former suffered high infection and the latter (which has a BCG program) did not. Survival prediction studies, using the BCG index as the predictor and controlling for other confounding factors such as age, gender, and environmental effects, could further explain the disparity of COVID-19 deaths between East and West. Finally, it is important to understand the precise virus-host interaction. Single-cell RNA sequencing in tissues with higher gene expression could be conducted to understand these dynamics. To bring the concept of precision epidemiology full circle, all the findings from bioinformatics analyses need to be further validated by experimental and clinical data.

This study conducts genetic probing with the intention of explaining the variability in symptoms and diverse outcomes of COVID-19. It provides some significant findings (SNP, CNV and gene expression) from *ACE2* and *TMPRSS2*, as evidence, for a plausible place to start looking into them. The work is a good first step to be followed by additional studies and functional assays that could potentially evaluate the findings to identify patients who may be at a higher risk of COVID-19-related mortality or infection, towards informed decisions for treatment and cure.

## List of abbreviations

COVID-19: 
*ACE2*: 
*TMPRSS2*: 
SNP: 
CNV: 
BCG: 
ExAC: 
TWCNV: 

## Ethics approval and consent to participate

Not applicable

## Consent for publication

Not Applicable

## Availability of data and material

Not applicable

## Authors’ contributions

E.Y.C., T.P.L., L.C.L. and M.H.T. has conceived the study. T.P.L. and A.C. co-wrote the manuscript. A.C. conducted all statistical analysis. C.Y.S. prepared the figures for the manuscript. E.Y.C., T.P.L., L.C.L. and M.H.T. has financially supported the work.

## Acknowledgements

We thank Melissa Stauffer, PhD, for editing the manuscript

## Supplementary Files

**Supplementary file 1.xlsx: Significant SNPs from gene ACE2.**

Description: List of 44 SNPs from ACE2 gene that are significant (P<0.05), across all sub-populations.

**Supplementary file 2.xlsx: Significant SNPs from gene TMPRSS2.**

Description: List of 98 SNPs from TMPRSS2 gene that are significant (P<0.05), across all sub-populations.

**Supplementary file 3.docx: Gene expression of tissues from healthy individuals.**

Description: Tissue specific gene expression in the GTEx healthy population

**Supplementary file 4.docx: Gene expression of patients with cancers in different tissues from CellExpress, stratified by gender.**

Description: The x-axis displays tissue names; the y-axis displays norm = log_2_ normalized gene expression values. Pink box plots display gene expression for females; blue box plots display gene expression for males. The upper panel shows gene expression for *ACE2*; the lower panel shows gene expression for *TMPRSS2*. All gene expression data were downloaded from CellExpress (http://cellexpress.cgm.ntu.edu.tw/).

**Supplementary file 5.docx: Gene expression of patients with cancers in different tissues from CellExpress, stratified by age.**

Description: The x-axis displays tissue names; the y-axis displays norm = log_2_ normalized gene expression values. Pink box-plots display gene expression for people with age <40 years; blue box-plots display gene expression for people with age >40 years and <60 years; and green box-plots display gene expression for people with age >60 years. The upper panel shows gene expression for *ACE2*; the lower panel shows gene expression for *TMPRSS2*. All gene expression data were downloaded from CellExpress (http://cellexpress.cgm.ntu.edu.tw/).

