## Supplementary file for "COVID-19: Variant screening, an important step towards precision epidemiology"

Eric Y. Chuang

Department of Electrical Engineering, Graduate Institute of Biomedical Electronics and Bioinformatics, National Taiwan University, Taipei 10617, Taiwan

**Supplementary file 1: Significant SNPs from Gene ACE2**

Gene: ACE2
Significance Threshold: p-value < 0.05
Comparison Done using ExAC subpopulations

2-way population comparison using Fisher's Exact Test for each SNP
p-val _overall: ad-hoc p-values for allele frequency over all populations

AFR-African
AMR: American (Latino)
EAS: East Asian
FIN: Finnish
NFE: Non-Finnish European
OTH: Others
SAS: South Asian

| chromosome | pos | ref | alt | rsID | AFR_AMR | AFR_EAS | AFR_FIN | AFR_NFE |
|---|---|---|---|---|---|---|---|---|
| X | 15582209 | C | T | rs35803318 | 8.51E-89 | 2.55E-11 | 0.001660281 | 1.03E-59 |
| X | 15582265 | G | A | rs147311723 | 2.21E-23 | 2.82E-23 | 4.49E-19 | 6.17E-74 |
| X | 15582298 | T | C | rs41303171 | 0.376166193 | 3.33E-05 | 4.84E-18 | 1.03E-32 |
| X | 15582359 | A | G | rs770208600 | 1 | 1 | 1 | 1 |
| X | 15584363 | A | C | rs764027290 | 0.127064363 | 1 | 1 | 1 |
| X | 15584367 | A | G | rs536749578 | 1 | 1 | 1 | 1 |
| X | 15584401 | T | C | rs751603885 | 1 | 1 | 1 | 1 |
| X | 15584416 | A | G | rs149039346 | 1.27E-06 | 2.38E-06 | 2.56E-05 | 1.40E-18 |
| X | 15584420 | A | G | rs4646179 | 3.21E-87 | 5.30E-92 | 6.39E-75 | 5.14E-264 |
| X | 15584515 | T | A | rs754997600 | 1 | 1 | 1 | 1 |
| X | 15589702 | G | A | rs4646169 | 2.21E-23 | 2.82E-23 | 4.49E-19 | 6.07E-71 |
| X | 15589725 | C | G | rs4646168 | 1.15E-175 | 6.64E-176 | 1.48E-143 | 0 |
| X | 15589737 | G | C | rs769144932 | 1 | 0.093565889 | 1 | 1 |
| X | 15589793 | G | T | rs145437639 | 0.106022834 | 0.256258957 | 0.287145014 | 0.002452983 |
| X | 15589821 | A | G | rs760929786 | 1 | 1 | 1 | 1 |
| X | 15589943 | G | C | rs766898296 | 0.252063543 | 1 | 1 | 1 |
| X | 15590284 | G | A | rs749275513 | 0.252063543 | 1 | 1 | 1 |
| X | 15591480 | T | C | rs759966064 | 1 | 1 | 1 | 1 |
| X | 15591485 | T | C | rs202137736 | 1 | 0.001799767 | 1 | 1 |
| X | 15593829 | T | C | rs191860450 | 1 | 3.85E-14 | 1 | 1 |
| X | 15593969 | G | A | rs754870498 | 1 | 1 | 1 | 1 |
| X | 15596345 | T | C | rs773676270 | 1 | 1 | 1 | 1 |
| X | 15599363 | G | A | rs147464721 | 2.96E-06 | 5.06E-08 | 1.44E-06 | 1.03E-23 |
| X | 15599392 | T | C | rs138390800 | 6.36E-07 | 1.24E-06 | 1.46E-05 | 1.96E-19 |
| X | 15603556 | G | A | rs781477897 | 1 | 1 | 1 | 1 |
| X | 15603587 | C | G | rs771528964 | 1 | 0.093565889 | 1 | 1 |
| X | 15603747 | A | G | rs768006201 | 0.224032098 | 0.504172638 | 0.524763342 | 0.018191504 |
| X | 15605847 | G | A | rs367784090 | 0.106022834 | 0.256258957 | 0.287145014 | 0.002452983 |
| X | 15605852 | C | T | rs4646140 | 4.26E-153 | 5.10E-77 | 2.51E-141 | 0 |
| X | 15607507 | C | T | rs759590772 | 1 | 1 | 1 | 1 |
| X | 15607532 | C | T | rs148771870 | 1 | 1 | 0.035901799 | 0.002449131 |
| X | 15609789 | G | A | rs772092723 | 0.224032098 | 0.504172638 | 0.524763342 | 0.018191504 |
| X | 15610328 | C | T | rs73195520 | 0.18509784 | 0.504172638 | 0.00019568 | 2.35E-05 |
| X | 15610348 | C | T | rs2285666 | 3.00E-99 | 4.11E-252 | 0.155294382 | 0.247564529 |
| X | 15610452 | A | T | rs528054982 | 1 | 1 | 1 | 1 |
| X | 15610461 | A | G | rs200477770 | 1 | 0.093565889 | 1 | 1 |
| X | 15613138 | G | A | rs757019762 | 1 | 0.206126193 | 1 | 1 |
| X | 15618828 | A | T | rs748232717 | 1 | 1 | 1 | 1 |
| X | 15618933 | G | A | rs368655410 | 1 | 0.093565889 | 7.79E-05 | 0.098365339 |
| X | 15618958 | T | C | rs4646116 | 0.614749362 | 0.135952959 | 1 | 6.86E-07 |
| X | 15618980 | A | G | rs73635825 | 1.40E-06 | 3.33E-05 | 0.000285172 | 2.13E-16 |
| X | 15618996 | A | G | rs779538833 | 1 | 0.093565889 | 1 | 1 |
| X | 15619055 | C | T | rs762634913 | 1 | 1 | 1 | 1 |
| X | 15619068 | T | C | rs370596467 | 1 | 0.000167887 | 1 | 1 |

| AFR_OTH | AFR_SAS | AMR_EAS | AMR_FIN | AMR_NFE | AMR_OTH | AMR_SAS | EAS_FIN | EAS_NFE |
|---|---|---|---|---|---|---|---|---|
| 6.03E-06 | 0.000390034 | 1.04E-119 | 7.62E-47 | 2.46E-21 | 0.000129402 | 2.35E-150 | 2.37E-19 | 1.28E-90 |
| 0.001096262 | 2.06E-36 | 0.140645709 | 0.303761879 | 0.002104 | 1 | 0.028844469 | 1 | 1 |
| 0.032708686 | 0.339079971 | 0.000535401 | 1.47E-22 | 4.36E-41 | 0.009847524 | 0.047070319 | 4.97E-30 | 9.51E-46 |
| 0.080254552 | 1 | 1 | 1 | 0.147833193 | 0.14016529 | 0.412175151 | 1 | 1 |
| 1 | 1 | 0.140645709 | 0.303761879 | 0.002104 | 1 | 0.028844469 | 1 | 1 |
| 1 | 0.004895784 | 1 | 1 | 1 | 1 | 0.002129829 | 1 | 1 |
| 0.080254552 | 0.000280219 | 1 | 1 | 1 | 0.072721448 | 0.000204228 | 1 | 1 |
| 0.252710889 | 8.08E-10 | 1 | 1 | 0.27381495 | 1 | 0.412175151 | 1 | 1 |
| 8.85E-12 | 2.58E-139 | 2.70E-05 | 0.000370793 | 3.58E-06 | 1 | 3.73E-06 | 1 | 0.103267798 |
| 0.080254552 | 1 | 1 | 1 | 1 | 0.072721448 | 1 | 1 | 1 |
| 0.001096262 | 2.06E-36 | 0.140645709 | 0.303761879 | 0.011478023 | 1 | 0.028844469 | 1 | 1 |
| 2.87E-19 | 6.90E-258 | 3.49E-07 | 6.91E-06 | 1.51E-13 | 0.280878553 | 1.91E-05 | 1 | 0.392989125 |
| 1 | 1 | 0.078227935 | 1 | 1 | 1 | 1 | 0.263292779 | 0.00151138 |
| 1 | 0.057752323 | 1 | 1 | 1 | 1 | 1 | 1 | 1 |
| 0.080254552 | 1 | 1 | 1 | 1 | 0.072721448 | 1 | 1 | 1 |
| 1 | 1 | 0.265592689 | 0.558145363 | 0.003229416 | 1 | 0.070002426 | 1 | 1 |
| 1 | 1 | 0.265592689 | 0.558145363 | 0.003229416 | 1 | 0.070002426 | 1 | 1 |
| 0.080254552 | 1 | 1 | 1 | 1 | 0.072721448 | 1 | 1 | 1 |
| 1 | 1 | 0.001116388 | 1 | 1 | 1 | 1 | 0.011857454 | 3.00E-08 |
| 1 | 1 | 3.74E-15 | 1 | 1 | 1 | 1 | 3.59E-10 | 6.61E-36 |
| 1 | 0.16390791 | 1 | 1 | 1 | 1 | 0.082033359 | 1 | 1 |
| 1 | 0.088322934 | 1 | 1 | 1 | 1 | 0.046121016 | 1 | 1 |
| 0.726216084 | 2.65E-12 | 0.140645709 | 0.303761879 | 0.002104 | 0.314517314 | 0.028844469 | 1 | 1 |
| 0.717193704 | 3.12E-10 | 1 | 1 | 0.27381495 | 0.14016529 | 0.412175151 | 1 | 1 |
| 1 | 0.288538428 | 1 | 1 | 1 | 1 | 0.273087856 | 1 | 1 |
| 1 | 1 | 0.078227935 | 1 | 1 | 1 | 1 | 0.263292779 | 0.00151138 |
| 1 | 0.149427612 | 1 | 1 | 1 | 1 | 1 | 1 | 1 |
| 1 | 0.057752323 | 1 | 1 | 1 | 1 | 1 | 1 | 1 |
| 2.24E-23 | 6.88E-26 | 4.35E-11 | 3.67E-09 | 1.40E-29 | 0.179433205 | 4.76E-79 | 3.87E-26 | 1.85E-86 |
| 1 | 0.088322934 | 1 | 1 | 1 | 1 | 0.046121016 | 1 | 1 |
| 1 | 0.000965722 | 1 | 0.026525622 | 0.001017282 | 1 | 0.000435632 | 0.015228788 | 0.000977766 |
| 1 | 0.149427612 | 1 | 1 | 1 | 1 | 1 | 1 | 1 |
| 0.221994193 | 0.149427612 | 0.022679694 | 0.010073367 | 0.004079807 | 0.453577174 | 0.002016722 | 1.87E-05 | 1.84E-06 |
| 0.00446946 | 1.26E-249 | 2.01E-49 | 1.85E-87 | 2.51E-202 | 6.04E-08 | 2.12E-31 | 3.48E-215 | 0 |
| 0.080254552 | 1 | 1 | 1 | 1 | 0.072721448 | 1 | 1 | 1 |
| 1 | 1 | 0.078227935 | 1 | 1 | 1 | 1 | 0.263292779 | 0.00151138 |
| 1 | 1 | 0.182935799 | 1 | 1 | 1 | 1 | 0.508861853 | 0.013172595 |
| 1 | 0.015214972 | 1 | 1 | 1 | 1 | 0.013103802 | 1 | 1 |
| 0.080254552 | 0.525692285 | 0.319766253 | 0.000294861 | 0.35050079 | 0.14016529 | 1 | 0.02169633 | 0.742367997 |
| 1 | 1 | 0.051270389 | 0.554167471 | 3.78E-06 | 1 | 0.644144347 | 0.322615125 | 1.60E-09 |
| 0.392399082 | 3.65E-08 | 1 | 1 | 1 | 1 | 1 | 1 | 1 |
| 1 | 1 | 0.078227935 | 1 | 1 | 1 | 1 | 0.263292779 | 0.00151138 |
| 1 | 0.088322934 | 1 | 1 | 1 | 1 | 0.046121016 | 1 | 1 |
| 1 | 0.015214972 | 8.71E-05 | 1 | 1 | 1 | 0.013103802 | 0.003470769 | 4.51E-11 |

| EAS_OTH | EAS_SAS | FIN_NFE | FIN_OTH | FIN_SAS | NFE_OTH | NFE_SAS | OTH_SAS | p-val_overall |
|---|---|---|---|---|---|---|---|---|
| 3.43E-17 | 1.62E-05 | 3.27E-24 | 0.00666865 | 1.85E-11 | 0.190234696 | 3.31E-123 | 1.65E-10 | 0.00049975 |
| 1 | 1 | 1 | 1 | 1 | 1 | 1 | 1 | 0.00049975 |
| 7.57E-06 | 1.31E-07 | 0.771053036 | 0.090382925 | 8.70E-19 | 0.049243846 | 6.16E-42 | 0.071258219 | 0.00049975 |
| 0.094959214 | 1 | 1 | 0.120712576 | 1 | 0.013422422 | 1 | 0.052123995 | 0.005497251 |
| 1 | 1 | 1 | 1 | 1 | 1 | 1 | 1 | 0.010994503 |
| 1 | 0.011230242 | 1 | 1 | 0.024403762 | 1 | 3.68E-09 | 1 | 0.00049975 |
| 0.094959214 | 0.001249377 | 1 | 0.120712576 | 0.005347178 | 0.013422422 | 1.12E-12 | 0.618839896 | 0.00049975 |
| 1 | 1 | 1 | 1 | 1 | 1 | 1 | 1 | 0.00049975 |
| 0.094959214 | 0.54871313 | 0.162882729 | 0.120712576 | 1 | 0.286771655 | 0.143633927 | 0.148379003 | 0.00049975 |
| 0.094959214 | 1 | 1 | 0.120712576 | 1 | 0.013422422 | 1 | 0.052123995 | 0.006996502 |
| 1 | 1 | 1 | 1 | 1 | 1 | 1 | 1 | 0.00049975 |
| 8.03E-05 | 0.05738266 | 0.630391413 | 0.000209869 | 0.115174422 | 8.44E-05 | 0.061450481 | 0.002579227 | 0.00049975 |
| 1 | 0.040645314 | 1 | 1 | 1 | 1 | 1 | 1 | 0.003998001 |
| 1 | 1 | 1 | 1 | 1 | 1 | 1 | 1 | 0.004997501 |
| 0.094959214 | 1 | 1 | 0.120712576 | 1 | 0.013422422 | 1 | 0.052123995 | 0.007496252 |
| 1 | 1 | 1 | 1 | 1 | 1 | 1 | 1 | 0.007996002 |
| 1 | 1 | 1 | 1 | 1 | 1 | 1 | 1 | 0.010494753 |
| 0.094959214 | 1 | 1 | 0.120712576 | 1 | 0.013422422 | 1 | 0.052123995 | 0.010994503 |
| 1 | 0.000194705 | 1 | 1 | 1 | 1 | 1 | 1 | 0.00049975 |
| 0.047518585 | 1.98E-17 | 1 | 1 | 1 | 1 | 0.35734118 | 1 | 0.00049975 |
| 1 | 0.17219569 | 1 | 1 | 0.330511148 | 1 | 0.000306622 | 1 | 0.003498251 |
| 1 | 0.100292209 | 1 | 1 | 0.192282297 | 1 | 0.000353202 | 1 | 0.006996502 |
| 0.094959214 | 1 | 1 | 0.120712576 | 1 | 0.026665074 | 1 | 0.052123995 | 0.00049975 |
| 0.094959214 | 1 | 1 | 0.120712576 | 1 | 0.026665074 | 1 | 0.052123995 | 0.00049975 |
| 1 | 0.555833426 | 1 | 1 | 0.562534993 | 1 | 0.00779989 | 1 | 0.046476762 |
| 1 | 0.040645314 | 1 | 1 | 1 | 1 | 1 | 1 | 0.004497751 |
| 1 | 1 | 1 | 1 | 1 | 1 | 1 | 1 | 0.018490755 |
| 1 | 1 | 1 | 1 | 1 | 1 | 1 | 1 | 0.006996502 |
| 0.0003897 | 1.88E-27 | 1 | 0.226881856 | 1.70E-83 | 0.183511132 | 0 | 1.35E-12 | 0.00049975 |
| 1 | 0.100292209 | 1 | 1 | 0.192282297 | 1 | 0.000353202 | 1 | 0.003998001 |
| 1 | 0.000421127 | 0.831556175 | 1 | 0.501577076 | 1 | 0.40366085 | 0.621734116 | 0.00049975 |
| 1 | 1 | 1 | 1 | 1 | 1 | 1 | 1 | 0.023488256 |
| 0.094959214 | 1 | 0.53279306 | 1 | 8.42E-08 | 1 | 3.11E-11 | 0.052123995 | 0.00049975 |
| 1.70E-29 | 2.40E-07 | 0.430904572 | 0.000703924 | 7.63E-202 | 0.000884652 | 0 | 1.37E-21 | 0.00049975 |
| 0.094959214 | 1 | 1 | 0.120712576 | 1 | 0.013422422 | 0.198337577 | 0.101536753 | 0.004497751 |
| 1 | 0.040645314 | 1 | 1 | 1 | 1 | 1 | 1 | 0.005997001 |
| 1 | 0.118233224 | 1 | 1 | 1 | 1 | 1 | 1 | 0.013993003 |
| 1 | 0.032512928 | 1 | 1 | 0.068031215 | 1 | 4.73E-07 | 1 | 0.00049975 |
| 0.329165498 | 0.347249064 | 0.000193987 | 1 | 0.000131375 | 0.247131091 | 0.287364445 | 0.148379003 | 0.00049975 |
| 1 | 0.111133078 | 1.71E-05 | 1 | 0.769165247 | 0.119779959 | 1.07E-09 | 1 | 0.00049975 |
| 1 | 1 | 1 | 1 | 1 | 1 | 1 | 1 | 0.00049975 |
| 1 | 0.120669411 | 1 | 1 | 1 | 1 | 0.198337577 | 1 | 0.004497751 |
| 1 | 0.100292209 | 1 | 1 | 0.192282297 | 1 | 6.08E-05 | 1 | 0.00049975 |
| 0.614440733 | 0.06072124 | 1 | 1 | 0.068031215 | 1 | 4.73E-07 | 1 | 0.00049975 |

**Supplementary file 2: Significant SNPs from Gene TMPRSS2**

Gene: TMPRSS2
Significance Threshold: p-value < 0.05
Comparison Done using ExAC subpopulations

2-way population comparison using Fisher's Exact Test for each SNP
p-val _overall: ad-hoc p-values for allele frequency over all populations

AFR-African
AMR: American (Latino)
EAS: East Asian
FIN: Finnish
NFE: Non-Finnish European
OTH: Others
SAS: South Asian

| chromosome | pos | ref | alt | rsID | AFR_AMR | AFR_EAS | AFR_FIN | AFR_NFE |
|---|---|---|---|---|---|---|---|---|
| 21 | 42838022 | C | T | rs200164183 | 1 | 0.042466558 | 1 | 1 |
| 21 | 42838036 | G | A | rs374981614 | 1 | 0.042466558 | 1 | 1 |
| 21 | 42838063 | T | C | rs370347248 | 0.016216241 | 0.009674312 | 0.026527209 | 8.64E-07 |
| 21 | 42838104 | G | C | rs118028230 | 0.378435142 | 9.91E-136 | 1 | 0.352539764 |
| 21 | 42838114 | T | C | rs369092228 | 0.000265783 | 0.001446336 | 0.009134339 | 2.67E-10 |
| 21 | 42839626 | G | A | rs183984610 | 0.04139043 | 1 | 1 | 0.011805018 |
| 21 | 42839661 | C | T | rs142750000 | 0.368793662 | 0.018399522 | 0.047926093 | 0.00440083 |
| 21 | 42839865 | G | A | rs147359020 | 0.104435677 | 0.067736469 | 0.415074765 | 0.011638597 |
| 21 | 42840494 | C | G | rs756459090 | 1 | 0.042466558 | 1 | 1 |
| 21 | 42842543 | C | T | rs78503214 | 1.58E-51 | 3.15E-50 | 8.62E-41 | 6.00E-158 |
| 21 | 42842570 | C | G | rs372285785 | 0.106022834 | 0.256258957 | 0.287145014 | 0.002452983 |
| 21 | 42842585 | G | A | rs146132415 | 0.085197313 | 0.03519287 | 0.088076356 | 3.72E-05 |
| 21 | 42842590 | C | T | rs369579311 | 0.224032098 | 0.504172638 | 0.524763342 | 0.018191504 |
| 21 | 42842591 | C | T | rs61735794 | 3.47E-05 | 6.94E-06 | 7.56E-20 | 1.81E-34 |
| 21 | 42842623 | G | A | rs61735795 | 1.33E-05 | 0.000120181 | 0.000814342 | 1.22E-12 |
| 21 | 42842717 | T | A | rs777860329 | 9.44E-09 | 1.97E-05 | 1.55E-10 | 8.01E-41 |
| 21 | 42843688 | T | C | rs370818663 | 0.224032098 | 0.504172638 | 0.524763342 | 0.018191504 |
| 21 | 42843704 | C | A | rs376752614 | 1 | 1 | 1 | 1 |
| 21 | 42843722 | G | A | rs376728899 | 0.002512957 | 0.046168614 | 0.026527209 | 1.09E-07 |
| 21 | 42843822 | T | C | rs565237319 | 1 | 1 | 1 | 1 |
| 21 | 42843843 | T | C | rs144046631 | 0.106022834 | 0.256258957 | 0.287145014 | 0.0088198 |
| 21 | 42843860 | T | C | rs768903594 | 0.016531386 | 1 | 1 | 1 |
| 21 | 42843873 | G | A | rs748528218 | 1 | 0.093565889 | 1 | 1 |
| 21 | 42843887 | A | T | rs769164587 | 1 | 1 | 1 | 1 |
| 21 | 42843935 | T | C | rs201320664 | 1 | 1 | 1 | 0.115843406 |
| 21 | 42843940 | G | A | rs187078345 | 1.07E-27 | 5.65E-25 | 2.57E-20 | 3.63E-79 |
| 21 | 42843951 | A | G | rs533556786 | 1 | 1 | 1 | 1 |
| 21 | 42845207 | G | A | rs75168613 | 3.42E-159 | 1.94E-181 | 1.52E-153 | 0 |
| 21 | 42845214 | C | A | rs577285910 | 1 | 0.093565889 | 1 | 1 |
| 21 | 42845216 | T | G | rs113288437 | 8.13E-174 | 4.49E-233 | 4.00E-196 | 0 |
| 21 | 42845220 | C | A | rs140141551 | 7.82E-12 | 6.49E-05 | 9.02E-06 | 2.48E-54 |
| 21 | 42845229 | G | A | rs201984814 | 0.032542788 | 1 | 1 | 1 |
| 21 | 42845239 | G | A | rs781723682 | 1 | 0.001799767 | 1 | 1 |
| 21 | 42845260 | G | A | rs143291395 | 1.40E-06 | 0.062600162 | 0.000285172 | 2.13E-16 |
| 21 | 42845329 | G | A | rs546335233 | 1 | 1 | 1 | 1 |
| 21 | 42845359 | C | T | rs2298658 | 4.19E-07 | 2.99E-07 | 0.388601645 | 1 |
| 21 | 42845366 | G | A | rs150445636 | 0.106022834 | 0.631234152 | 0.287145014 | 0.0088198 |
| 21 | 42845373 | C | T | rs61735796 | 1 | 1 | 1 | 0.063355608 |
| 21 | 42845374 | G | A | rs2298659 | 2.17E-24 | 7.45E-18 | 4.93E-84 | 4.09E-08 |
| 21 | 42845383 | A | G | rs17854725 | 0.83088908 | 5.53E-178 | 6.97E-36 | 3.58E-91 |
| 21 | 42845385 | T | C | rs745679574 | 1 | 1 | 1 | 1 |
| 21 | 42845438 | C | T | rs372744084 | 0.050170055 | 0.131267226 | 0.162448855 | 0.000330711 |
| 21 | 42845459 | C | T | rs368812287 | 0.00118853 | 0.005112524 | 0.014955756 | 6.24E-07 |
| 21 | 42845470 | C | G | rs777296694 | 0.501375696 | 1 | 1 | 1 |
| 21 | 42845473 | C | G | rs199704563 | 0.47334425 | 1 | 1 | 0.240750803 |
| 21 | 42848457 | C | T | rs75756279 | 0.220163457 | 0.067736469 | 0.000299167 | 2.70E-08 |
| 21 | 42848494 | A | C | rs200615061 | 1 | 1 | 1 | 0.580810495 |
| 21 | 42848536 | G | A | rs547544037 | 1 | 1 | 1 | 1 |
| 21 | 42848546 | C | A | rs143597099 | 1 | 1 | 0.008845446 | 1 |
| 21 | 42848563 | G | A | rs754886975 | 0.00118853 | 0.005112524 | 0.014955756 | 1.47E-08 |
| 21 | 42848571 | C | G | rs199999942 | 0.004296395 | 1 | 1 | 1 |

| | | | | | | | | |
|---|---|---|---|---|---|---|---|---|
| 21 | 42848579 | G | A | rs74423429 | 0.886308307 | 6.94E-06 | 2.73E-25 | 2.26E-31 |
| 21 | 42848591 | T | G | rs375891843 | 1 | 1 | 0.388601645 | 0.097755222 |
| 21 | 42848593 | A | C | rs765703243 | 1.64E-11 | 6.26E-09 | 5.14E-05 | 1.01E-05 |
| 21 | 42848596 | T | G | rs775850899 | 0.224032098 | 0.504172638 | 0.524763342 | 0.018191504 |
| 21 | 42851134 | G | A | rs141788162 | 0.54396155 | 0.695558162 | 1.64E-07 | 0.142235716 |
| 21 | 42851157 | A | T | rs150554820 | 0.127064363 | 1 | 1 | 0.065519757 |
| 21 | 42851197 | G | A | rs146681599 | 0.224032098 | 0.504172638 | 0.524763342 | 0.018191504 |
| 21 | 42851239 | G | T | rs751618573 | 1 | 1 | 1 | 1 |
| 21 | 42851258 | A | G | rs377385043 | 1 | 1 | 0.058650665 | 0.001318788 |
| 21 | 42852365 | C | T | rs186734573 | 0.350475477 | 0.256258957 | 2.38E-13 | 2.41E-07 |
| 21 | 42852382 | C | A | rs79397218 | 1 | 1 | 1 | 1 |
| 21 | 42852435 | G | A | rs61735789 | 2.03E-05 | 0.000773433 | 0.03521051 | 6.84E-16 |
| 21 | 42852497 | C | T | rs12329760 | 1.28E-65 | 6.68E-21 | 1.07E-15 | 5.03E-20 |
| 21 | 42852536 | G | A | rs147465180 | 1 | 1 | 1 | 1 |
| 21 | 42852552 | C | T | rs374531004 | 0.057956613 | 0.03519287 | 0.088076356 | 6.01E-06 |
| 21 | 42860333 | C | T | rs758778273 | 1 | 0.093565889 | 1 | 1 |
| 21 | 42860346 | G | A | rs768354120 | 0.008417954 | 1 | 1 | 1 |
| 21 | 42860488 | A | G | rs187290362 | 0.06942605 | 0.504172638 | 1 | 0.002285423 |
| 21 | 42861434 | T | C | rs368936645 | 0.224032098 | 0.504172638 | 0.524763342 | 0.018191504 |
| 21 | 42861455 | C | A | rs781008294 | 1 | 1 | 1 | 1 |
| 21 | 42861476 | G | A | rs190265904 | 1 | 0.093565889 | 1 | 1 |
| 21 | 42861524 | G | C | . | 1 | 1 | 1 | 1 |
| 21 | 42861545 | G | A | rs144948620 | 0.129837182 | 0.005112524 | 3.02E-16 | 3.19E-19 |
| 21 | 42861560 | G | A | rs574683527 | 1 | 1 | 1 | 1 |
| 21 | 42866232 | G | A | rs373196115 | 0.001135157 | 0.708828511 | 0.441893491 | 0.010757159 |
| 21 | 42866296 | T | C | rs3787950 | 4.60E-146 | 6.03E-09 | 1.84E-128 | 2.61E-134 |
| 21 | 42866297 | G | A | rs61735793 | 1 | 0.002713931 | 3.81E-07 | 4.19E-11 |
| 21 | 42866301 | C | G | rs114363287 | 0.005312709 | 0.018399522 | 0.047926093 | 8.10E-07 |
| 21 | 42866326 | G | A | rs778999405 | 1 | 1 | 1 | 1 |
| 21 | 42866329 | G | A | rs372804318 | 0.47334425 | 1 | 1 | 0.348825589 |
| 21 | 42866332 | G | A | rs61735792 | 2.81E-08 | 0.000829993 | 3.37E-12 | 2.65E-13 |
| 21 | 42866377 | G | A | rs149527323 | 0.000125667 | 0.000773433 | 0.004874291 | 2.50E-09 |
| 21 | 42866388 | A | C | rs201679623 | 1 | 5.28E-10 | 1 | 1 |
| 21 | 42866396 | G | A | rs376143876 | 1 | 1 | 1 | 1 |
| 21 | 42866410 | G | A | rs187460831 | 0.47334425 | 0.098355735 | 1 | 0.251582691 |
| 21 | 42866422 | G | A | rs199824558 | 0.350499515 | 0.256258957 | 0.287145014 | 1 |
| 21 | 42866423 | A | G | rs201093031 | 1 | 0.000167887 | 1 | 1 |
| 21 | 42866439 | C | T | rs61735791 | 1 | 0.398308523 | 0.415074765 | 0.069194308 |
| 21 | 42866468 | T | C | rs61735790 | 1.50E-11 | 3.56E-12 | 4.62E-10 | 1.75E-37 |
| 21 | 42866553 | T | C | rs368773451 | 0.378435142 | 1 | 5.38E-15 | 7.58E-07 |
| 21 | 42866595 | C | A | rs745378739 | 1 | 0.093565889 | 1 | 1 |
| 21 | 42866638 | A | G | rs143523726 | 0.011230598 | 0.03519287 | 0.088076356 | 6.01E-06 |
| 21 | 42866644 | G | A | rs368625065 | 1 | 1 | 1 | 0.001984052 |
| 21 | 42866689 | T | C | rs774042268 | 0.023738088 | 0.067736469 | 0.163882047 | 4.46E-05 |
| 21 | 42879865 | G | C | rs562751769 | 1 | 1 | 1 | 1 |
| 21 | 42879909 | C | A | rs75603675 | 0.020646344 | 0 | 1 | 2.17E-14 |

| AFR_OTH | AFR_SAS | AMR_EAS | AMR_FIN | AMR_NFE | AMR_OTH | AMR_SAS | EAS_FIN | EAS_NFE |
|---|---|---|---|---|---|---|---|---|
| 1 | 1 | 0.033447799 | 1 | 1 | 1 | 1 | 0.1383309 | 0.000173375 |
| 1 | 1 | 0.033447799 | 1 | 1 | 1 | 1 | 0.1383309 | 0.000173375 |
| 1 | 0.000497157 | 1 | 1 | 0.27381495 | 1 | 0.412175151 | 1 | 1 |
| 0.001932115 | 0.060122884 | 3.01E-140 | 0.303761879 | 0.005570093 | 0.01071842 | 0.422071679 | 1.05E-99 | 0 |
| 1 | 2.86E-05 | 1 | 1 | 1 | 1 | 1 | 1 | 1 |
| 1 | 0.38658147 | 0.01259419 | 0.169105262 | 1 | 1 | 0.000830652 | 0.433193608 | 0.003378911 |
| 1 | 0.052909122 | 0.140645709 | 0.303761879 | 2.71E-05 | 1 | 0.456884839 | 1 | 5.59E-07 |
| 0.021190753 | 1.69E-67 | 0.000994645 | 0.024209177 | 0.497882546 | 0.105530667 | 3.20E-64 | 0.433193608 | 1.47E-05 |
| 1 | 1 | 0.033447799 | 1 | 1 | 1 | 1 | 0.1383309 | 0.000173375 |
| 3.57E-06 | 2.14E-78 | 0.022679694 | 0.053339399 | 0.000122017 | 0.453577174 | 0.002016722 | 1 | 1 |
| 1 | 0.057752323 | 1 | 1 | 1 | 1 | 1 | 1 | 1 |
| 1 | 0.003331791 | 0.000158898 | 0.001032373 | 7.49E-13 | 0.624675695 | 6.86E-07 | 1 | 1 |
| 1 | 0.149427612 | 1 | 1 | 1 | 1 | 1 | 1 | 1 |
| 6.05E-07 | 2.42E-05 | 1.10E-15 | 1.32E-08 | 2.18E-16 | 0.002118965 | 0.872563074 | 7.80E-34 | 4.09E-50 |
| 0.626428692 | 6.36E-07 | 1 | 1 | 1 | 1 | 1 | 1 | 1 |
| 1 | 9.12E-20 | 0.341568531 | 0.057583302 | 2.27E-07 | 1 | 0.000830652 | 0.006667771 | 3.93E-10 |
| 1 | 0.149427612 | 1 | 1 | 1 | 1 | 1 | 1 | 1 |
| 1 | 2.91E-07 | 1 | 1 | 1 | 1 | 8.45E-08 | 1 | 1 |
| 1 | 0.000497157 | 0.427738236 | 1 | 1 | 1 | 1 | 1 | 0.11478367 |
| 1 | 0.04824904 | 1 | 1 | 1 | 1 | 0.046396601 | 1 | 1 |
| 1 | 0.057752323 | 1 | 1 | 1 | 1 | 1 | 1 | 1 |
| 0.080254552 | 1 | 0.022679694 | 0.053339399 | 0.000122017 | 0.453577174 | 0.002016722 | 1 | 1 |
| 1 | 1 | 0.078227935 | 1 | 1 | 1 | 1 | 0.263292779 | 0.00151138 |
| 1 | 0.288538428 | 1 | 1 | 1 | 1 | 0.273087856 | 1 | 1 |
| 1 | 0.38658147 | 1 | 1 | 0.07712139 | 1 | 0.412175151 | 1 | 0.04369821 |
| 0.006285145 | 6.46E-39 | 0.510395126 | 0.537177371 | 0.059095474 | 0.202713753 | 0.169871103 | 1 | 1 |
| 0.080254552 | 1.67E-11 | 1 | 1 | 1 | 0.072721448 | 1.92E-12 | 1 | 1 |
| 1.48E-12 | 1.28E-213 | 3.12E-11 | 7.18E-12 | 1.66E-08 | 0.000562923 | 0.114218626 | 0.263292779 | 0.000173271 |
| 1 | 1 | 0.078227935 | 1 | 1 | 1 | 1 | 0.263292779 | 0.00151138 |
| 1.26E-17 | 2.28E-275 | 1.44E-23 | 8.83E-23 | 1.52E-27 | 0.05588337 | 2.74E-07 | 0.263292779 | 2.02E-05 |
| 2.81E-08 | 0.111301023 | 1.39E-22 | 0.116764616 | 1.47E-18 | 0.021116733 | 7.56E-10 | 1.38E-14 | 1.87E-65 |
| 1 | 0.008780307 | 0.041194522 | 0.092916628 | 0.000222558 | 1 | 0.80614737 | 1 | 1 |
| 1 | 1 | 0.001116388 | 1 | 1 | 1 | 1 | 0.011857454 | 3.00E-08 |
| 0.392399082 | 3.65E-08 | 0.006112324 | 1 | 1 | 1 | 1 | 0.0397011 | 2.28E-06 |
| 1 | 0.163866662 | 1 | 1 | 1 | 1 | 0.148205248 | 1 | 1 |
| 1 | 1 | 0.756808581 | 0.000413624 | 7.39E-16 | 0.407472435 | 1.99E-08 | 0.000342571 | 7.41E-15 |
| 1 | 0.381306408 | 0.427738236 | 1 | 1 | 1 | 0.515391904 | 1 | 0.216394744 |
| 1 | 1 | 1 | 1 | 0.039158947 | 1 | 1 | 1 | 0.101649575 |
| 0.001610087 | 0.302163246 | 0.377160954 | 2.42E-27 | 1.52E-15 | 0.422903818 | 4.98E-25 | 4.33E-28 | 3.44E-09 |
| 2.39E-05 | 5.54E-19 | 6.10E-183 | 2.40E-38 | 1.52E-102 | 1.31E-05 | 3.93E-21 | 1.06E-304 | 0 |
| 0.080254552 | 1 | 1 | 1 | 1 | 0.072721448 | 1 | 1 | 1 |
| 1 | 0.02231808 | 1 | 1 | 1 | 1 | 1 | 1 | 1 |
| 1 | 0.00019201 | 1 | 1 | 1 | 1 | 1 | 1 | 1 |
| 1 | 1 | 0.510395126 | 0.537177371 | 0.021851436 | 1 | 0.169871103 | 1 | 1 |
| 0.154081361 | 0.38658147 | 1 | 1 | 0.039158947 | 0.072721448 | 1 | 1 | 0.101649575 |
| 0.021190753 | 4.29E-07 | 0.00351791 | 0.009806649 | 8.45E-06 | 0.076284895 | 5.76E-05 | 6.51E-07 | 1.69E-11 |
| 0.154081361 | 3.96E-05 | 0.510395126 | 0.537177371 | 0.277360312 | 0.202713753 | 0.000133658 | 1 | 1 |
| 1 | 0.026837664 | 1 | 1 | 1 | 1 | 0.024554828 | 1 | 1 |
| 0.080254552 | 1 | 1 | 0.0063399 | 1 | 0.072721448 | 1 | 0.015228788 | 1 |
| 1 | 0.00019201 | 1 | 1 | 1 | 1 | 1 | 1 | 1 |
| 1 | 1 | 0.012795782 | 0.031154728 | 3.35E-08 | 1 | 0.000342096 | 1 | 1 |

| | | | | | | | | |
|---|---|---|---|---|---|---|---|---|
| 0.003037087 | 0.017108323 | 1.17E-05 | 1.29E-27 | 1.08E-35 | 0.002016065 | 0.005647487 | 1.05E-39 | 5.49E-47 |
| 0.006427743 | 1 | 1 | 0.363566403 | 0.098300879 | 0.005277606 | 1 | 0.433193608 | 0.157026627 |
| 0.606798577 | 1.05E-18 | 0.730941505 | 2.76E-24 | 6.34E-06 | 0.00019616 | 0.16683499 | 1.63E-19 | 0.000442844 |
| 1 | 0.149427612 | 1 | 1 | 1 | 1 | 1 | 1 | 1 |
| 0.131819372 | 0.001982609 | 0.303501352 | 2.16E-09 | 0.013095188 | 0.076284895 | 0.016054793 | 3.96E-06 | 0.432418032 |
| 0.080254552 | 1 | 0.140645709 | 0.303761879 | 1 | 0.314517314 | 0.028844469 | 1 | 0.103267798 |
| 1 | 0.149427612 | 1 | 1 | 1 | 1 | 1 | 1 | 1 |
| 1 | 0.163866662 | 1 | 1 | 1 | 1 | 0.148205248 | 1 | 1 |
| 1 | 1 | 1 | 0.048028653 | 0.000503666 | 1 | 1 | 0.08124988 | 0.003378911 |
| 0.008545989 | 0.305603635 | 1 | 1.64E-16 | 2.41E-10 | 0.00144603 | 1 | 5.96E-15 | 5.74E-09 |
| 0.080254552 | 9.59E-14 | 1 | 1 | 1 | 0.072721448 | 5.16E-15 | 1 | 1 |
| 0.113533834 | 0.026039892 | 6.29E-12 | 0.117590782 | 0.000560453 | 1 | 1.32E-12 | 6.51E-07 | 2.59E-24 |
| 0.517480545 | 3.74E-09 | 1.89E-147 | 2.70E-120 | 4.66E-40 | 6.29E-10 | 2.66E-38 | 0.535993876 | 4.54E-95 |
| 0.080254552 | 1 | 1 | 1 | 1 | 0.072721448 | 1 | 1 | 1 |
| 1 | 0.003331791 | 1 | 1 | 0.147833193 | 1 | 0.412175151 | 1 | 1 |
| 1 | 1 | 0.078227935 | 1 | 1 | 1 | 1 | 0.263292779 | 0.00151138 |
| 1 | 1 | 0.01259419 | 0.057583302 | 2.27E-07 | 1 | 0.000830652 | 1 | 1 |
| 0.000539016 | 0.09585713 | 0.012795782 | 0.105283462 | 0.357525613 | 0.011633464 | 1 | 0.433193608 | 0.000185924 |
| 1 | 0.149427612 | 1 | 1 | 1 | 1 | 1 | 1 | 1 |
| 0.080254552 | 1 | 1 | 1 | 1 | 0.072721448 | 1 | 1 | 1 |
| 0.080254552 | 1 | 0.319766253 | 1 | 0.27381495 | 0.14016529 | 1 | 0.263292779 | 0.005525395 |
| 0.080254552 | 1 | 1 | 1 | 1 | 0.072721448 | 1 | 1 | 1 |
| 0.000123588 | 0.562892674 | 2.70E-05 | 1.88E-12 | 9.61E-15 | 0.002002055 | 0.322329916 | 2.73E-22 | 3.05E-25 |
| 1 | 0.002668953 | 1 | 1 | 1 | 1 | 0.001189109 | 1 | 1 |
| 0.0546038 | 0.272888157 | 0.011633452 | 0.054444729 | 0.077921323 | 0.672044633 | 0.010096401 | 0.733557708 | 0.104435117 |
| 2.80E-06 | 0.608416446 | 3.91E-79 | 0.002510572 | 1.08E-23 | 1.25E-08 | 2.80E-167 | 2.97E-77 | 5.93E-50 |
| 0.021427552 | 0.021885351 | 0.001863142 | 2.52E-07 | 1.13E-11 | 0.024894069 | 0.026653318 | 4.03E-13 | 5.45E-18 |
| 1 | 0.001287097 | 1 | 1 | 1 | 1 | 1 | 1 | 1 |
| 0.080254552 | 1 | 1 | 1 | 1 | 0.072721448 | 1 | 1 | 1 |
| 1 | 0.38658147 | 1 | 1 | 0.061416103 | 1 | 1 | 1 | 0.161806811 |
| 8.13E-05 | 0.876993847 | 1.89E-16 | 0.034564317 | 0.533411269 | 0.187481232 | 3.89E-10 | 9.47E-21 | 1.18E-22 |
| 0.615876011 | 9.25E-05 | 1 | 1 | 1 | 1 | 1 | 1 | 1 |
| 1 | 1 | 1.73E-09 | 1 | 0.147833193 | 1 | 1 | 3.07E-07 | 3.85E-26 |
| 1 | 0.008780307 | 1 | 1 | 1 | 1 | 0.007067856 | 1 | 1 |
| 1 | 0.38658147 | 0.014299331 | 1 | 1 | 1 | 1 | 0.073672619 | 0.000107906 |
| 0.28446583 | 3.55E-12 | 0.022679694 | 0.053339399 | 0.093771661 | 0.453577174 | 2.21E-10 | 1 | 0.157026627 |
| 1 | 1 | 8.71E-05 | 1 | 1 | 1 | 1 | 0.003470769 | 4.51E-11 |
| 0.394791063 | 0.321076601 | 0.382468352 | 0.426787967 | 0.027040809 | 0.364406564 | 0.501962779 | 0.149241461 | 0.489894938 |
| 0.046771723 | 6.17E-19 | 0.265592689 | 0.558145363 | 0.011486053 | 1 | 0.070002426 | 1 | 1 |
| 0.154081361 | 1 | 0.140645709 | 3.46E-13 | 3.88E-05 | 0.314517314 | 0.16683499 | 1.38E-14 | 8.24E-07 |
| 1 | 1 | 0.078227935 | 1 | 1 | 1 | 1 | 0.263292779 | 0.00151138 |
| 1 | 0.003331791 | 1 | 1 | 1 | 1 | 1 | 1 | 1 |
| 1 | 1 | 1 | 1 | 0.001232698 | 1 | 1 | 1 | 0.008400531 |
| 1 | 0.008623686 | 1 | 1 | 1 | 1 | 1 | 1 | 1 |
| 1 | 0.04824904 | 1 | 1 | 1 | 1 | 0.046396601 | 1 | 1 |
| 0.652355998 | 8.68E-15 | 0 | 1 | 1.11E-06 | 0.166376205 | 2.88E-26 | 1 | 0 |

| EAS_OTH | EAS_SAS | FIN_NFE | FIN_OTH | FIN_SAS | NFE_OTH | NFE_SAS | OTH_SAS | p-val_overall |
|---|---|---|---|---|---|---|---|---|
| 1 | 0.013970615 | 1 | 1 | 1 | 1 | 1 | 1 | 0.0009995 |
| 1 | 0.013970615 | 1 | 1 | 1 | 1 | 1 | 1 | 0.00049975 |
| 1 | 1 | 1 | 1 | 1 | 1 | 1 | 1 | 0.00049975 |
| 5.00E-14 | 4.69E-168 | 1 | 0.001748755 | 0.072140653 | 2.35E-05 | 4.16E-06 | 0.027213391 | 0.00049975 |
| 1 | 1 | 1 | 1 | 1 | 1 | 1 | 1 | 0.00049975 |
| 1 | 1 | 0.085831627 | 1 | 0.285998443 | 1 | 2.27E-05 | 1 | 0.00049975 |
| 1 | 0.555833426 | 1.24E-05 | 1 | 0.562534993 | 0.419245584 | 8.01E-09 | 1 | 0.00049975 |
| 0.000851155 | 1.11E-65 | 0.00204362 | 0.006365413 | 1.30E-50 | 0.145129961 | 3.92E-159 | 1.27E-05 | 0.00049975 |
| 1 | 0.013970615 | 1 | 1 | 1 | 1 | 1 | 1 | 0.0009995 |
| 0.094959214 | 1 | 1 | 0.120712576 | 1 | 0.052620646 | 1 | 0.052123995 | 0.00049975 |
| 1 | 1 | 1 | 1 | 1 | 1 | 1 | 1 | 0.007496252 |
| 1 | 1 | 1 | 1 | 1 | 1 | 1 | 1 | 0.00049975 |
| 1 | 1 | 1 | 1 | 1 | 1 | 1 | 1 | 0.019490255 |
| 5.53E-14 | 3.28E-16 | 1 | 0.765692577 | 3.09E-10 | 0.776757336 | 8.22E-23 | 0.001382886 | 0.00049975 |
| 1 | 1 | 1 | 1 | 1 | 1 | 1 | 1 | 0.00049975 |
| 1 | 2.30E-05 | 1 | 1 | 1 | 1 | 1 | 1 | 0.00049975 |
| 1 | 1 | 1 | 1 | 1 | 1 | 1 | 1 | 0.022988506 |
| 1 | 3.24E-06 | 1 | 1 | 6.28E-05 | 1 | 1.58E-22 | 0.4074371 | 0.00049975 |
| 1 | 0.343876659 | 1 | 1 | 1 | 1 | 1 | 1 | 0.00049975 |
| 1 | 0.1038879 | 1 | 1 | 0.2030777 | 1 | 1.20E-05 | 1 | 0.00049975 |
| 1 | 1 | 1 | 1 | 1 | 1 | 1 | 1 | 0.053973013 |
| 0.094959214 | 1 | 1 | 0.120712576 | 1 | 0.052620646 | 1 | 0.052123995 | 0.00049975 |
| 1 | 0.040645314 | 1 | 1 | 1 | 1 | 1 | 1 | 0.002498751 |
| 1 | 0.555833426 | 1 | 1 | 0.562534993 | 1 | 0.00779989 | 1 | 0.047476262 |
| 1 | 1 | 0.108930985 | 1 | 1 | 1 | 0.002281793 | 1 | 0.00149925 |
| 0.094959214 | 1 | 1 | 0.120712576 | 1 | 0.026665074 | 1 | 0.052123995 | 0.00049975 |
| 0.094959214 | 5.80E-10 | 1 | 0.120712576 | 5.11E-08 | 0.026665074 | 5.45E-35 | 0.523847645 | 0.00049975 |
| 2.09E-12 | 1.04E-08 | 8.28E-06 | 1.17E-13 | 1.69E-09 | 8.49E-09 | 3.34E-05 | 3.13E-05 | 0.00049975 |
| 1 | 0.040645314 | 1 | 1 | 1 | 1 | 1 | 1 | 0.003498251 |
| 2.32E-13 | 7.25E-11 | 1.12E-06 | 1.37E-14 | 1.76E-11 | 8.98E-09 | 2.41E-06 | 5.59E-05 | 0.00049975 |
| 4.04E-15 | 1.54E-08 | 3.83E-17 | 0.003783007 | 0.000476066 | 0.702594015 | 1.11E-67 | 8.72E-07 | 0.00049975 |
| 1 | 0.018997025 | 1 | 1 | 0.072140653 | 1 | 4.16E-06 | 1 | 0.00049975 |
| 1 | 0.000194705 | 1 | 1 | 1 | 1 | 1 | 1 | 0.00049975 |
| 1 | 0.001649787 | 1 | 1 | 1 | 1 | 1 | 1 | 0.00049975 |
| 1 | 0.305951479 | 1 | 1 | 0.58352524 | 1 | 0.006506315 | 1 | 0.04197901 |
| 0.248698758 | 2.04E-08 | 0.376551949 | 1 | 0.490219439 | 1 | 1 | 1 | 0.00049975 |
| 1 | 1 | 1 | 1 | 1 | 1 | 0.102402144 | 1 | 0.023488256 |
| 1 | 1 | 0.261574014 | 1 | 1 | 1 | 0.014583339 | 1 | 0.007996002 |
| 0.662885458 | 2.26E-17 | 2.15E-84 | 5.00E-08 | 6.18E-91 | 0.125126569 | 2.28E-07 | 0.00445564 | 0.00049975 |
| 5.54E-64 | 0 | 0.192184306 | 0.159602439 | 4.43E-09 | 0.045583359 | 5.75E-32 | 0.311398873 | 0.00049975 |
| 0.094959214 | 1 | 1 | 0.120712576 | 1 | 0.013422422 | 1 | 0.052123995 | 0.008995502 |
| 1 | 1 | 1 | 1 | 1 | 1 | 1 | 1 | 0.002998501 |
| 1 | 1 | 1 | 1 | 1 | 1 | 1 | 1 | 0.00049975 |
| 1 | 1 | 1 | 1 | 1 | 1 | 1 | 1 | 0.048975512 |
| 0.094959214 | 1 | 0.261574014 | 0.120712576 | 1 | 0.277061175 | 0.014583339 | 0.052123995 | 0.004997501 |
| 0.000851155 | 3.67E-10 | 0.558350301 | 0.726154239 | 0.507380141 | 0.762759349 | 0.815724685 | 0.767334075 | 0.00049975 |
| 0.094959214 | 2.85E-05 | 1 | 0.120712576 | 0.000260408 | 0.077885554 | 3.09E-14 | 1 | 0.00049975 |
| 1 | 0.05738266 | 1 | 1 | 0.115174422 | 1 | 2.39E-06 | 1 | 0.00049975 |
| 0.094959214 | 1 | 3.30E-05 | 0.538100884 | 0.001909326 | 0.026665074 | 1 | 0.052123995 | 0.00049975 |
| 1 | 1 | 1 | 1 | 1 | 1 | 1 | 1 | 0.00049975 |
| 1 | 1 | 1 | 1 | 1 | 1 | 1 | 1 | 0.00049975 |

| | | | | | | | | |
|---|---|---|---|---|---|---|---|---|
| 5.16E-08 | 2.04E-11 | 0.034511831 | 0.084351897 | 4.54E-21 | 0.303866986 | 3.54E-30 | 0.058398977 | 0.00049975 |
| 0.008999273 | 1 | 1 | 0.040132381 | 0.285998443 | 0.03190314 | 0.021250772 | 0.002711238 | 0.002998501 |
| 0.000688889 | 0.050648061 | 1.44E-20 | 0.265300136 | 1.40E-34 | 0.041983382 | 3.50E-12 | 2.16E-06 | 0.00049975 |
| 1 | 1 | 1 | 1 | 1 | 1 | 1 | 1 | 0.022988506 |
| 0.189293062 | 0.000479732 | 4.95E-08 | 0.473577412 | 1.81E-17 | 0.432889759 | 8.17E-09 | 0.004206296 | 0.00049975 |
| 0.094959214 | 1 | 0.162882729 | 0.120712576 | 1 | 0.286771655 | 0.008484443 | 0.052123995 | 0.004997501 |
| 1 | 1 | 1 | 1 | 1 | 1 | 1 | 1 | 0.024487756 |
| 1 | 0.305951479 | 1 | 1 | 0.58352524 | 1 | 0.001546561 | 1 | 0.006496752 |
| 1 | 1 | 0.627669274 | 1 | 0.023378122 | 1 | 2.27E-05 | 1 | 0.00049975 |
| 0.000851155 | 1 | 4.68E-06 | 0.473577412 | 1.55E-20 | 0.48655165 | 1.78E-14 | 0.000541053 | 0.00049975 |
| 0.094959214 | 8.77E-12 | 1 | 0.120712576 | 1.69E-09 | 0.026665074 | 1.16E-42 | 0.261035068 | 0.00049975 |
| 0.000851155 | 0.100292209 | 9.15E-06 | 0.726154239 | 8.61E-06 | 0.298761756 | 5.04E-35 | 0.00933498 | 0.00049975 |
| 6.62E-06 | 5.32E-57 | 8.78E-69 | 3.00E-05 | 1.85E-43 | 0.033260769 | 0.011090139 | 0.145874615 | 0.00049975 |
| 0.094959214 | 1 | 1 | 0.120712576 | 1 | 0.026665074 | 1 | 0.052123995 | 0.047976012 |
| 1 | 1 | 1 | 1 | 1 | 1 | 1 | 1 | 0.00049975 |
| 1 | 0.040645314 | 1 | 1 | 1 | 1 | 1 | 1 | 0.005997001 |
| 1 | 1 | 1 | 1 | 1 | 1 | 1 | 1 | 0.00049975 |
| 8.03E-05 | 0.011230242 | 0.014372337 | 0.000948797 | 0.126668329 | 0.022778358 | 0.179751275 | 0.008032669 | 0.00049975 |
| 1 | 1 | 1 | 1 | 1 | 1 | 1 | 1 | 0.023988006 |
| 0.094959214 | 1 | 1 | 0.120712576 | 1 | 0.013422422 | 1 | 0.052123995 | 0.009495252 |
| 0.329165498 | 0.120669411 | 1 | 0.120712576 | 1 | 0.026665074 | 0.35734118 | 0.101536753 | 0.003498251 |
| 0.094959214 | 1 | 1 | 0.120712576 | 1 | 0.013422422 | 0.198337577 | 0.101536753 | 0.005497251 |
| 6.68E-08 | 0.000421127 | 0.076851956 | 0.849807433 | 8.09E-18 | 0.686968812 | 7.04E-25 | 0.00027617 | 0.00049975 |
| 1 | 0.00631156 | 1 | 1 | 0.025948718 | 1 | 7.29E-10 | 1 | 0.00049975 |
| 0.104380159 | 0.599761497 | 0.318789486 | 0.157055097 | 1 | 0.266707323 | 0.132754911 | 0.144831939 | 0.003998001 |
| 0.022060837 | 4.61E-09 | 1.37E-28 | 2.39E-12 | 3.10E-139 | 0.014421729 | 2.11E-182 | 5.04E-06 | 0.00049975 |
| 8.03E-05 | 3.73E-07 | 0.782959706 | 1 | 0.000457151 | 1 | 3.17E-07 | 0.153393812 | 0.00049975 |
| 1 | 1 | 1 | 1 | 1 | 1 | 1 | 1 | 0.00049975 |
| 0.094959214 | 1 | 1 | 0.120712576 | 1 | 0.013422422 | 0.198337577 | 0.101536753 | 0.005497251 |
| 1 | 1 | 0.25230071 | 1 | 1 | 1 | 0.01343437 | 1 | 0.036481759 |
| 5.35E-09 | 0.000185606 | 0.042140207 | 0.708184437 | 1.44E-14 | 0.218152696 | 1.46E-18 | 6.99E-05 | 0.00049975 |
| 1 | 1 | 1 | 1 | 1 | 1 | 0.484810698 | 1 | 0.00049975 |
| 0.104557449 | 5.40E-12 | 1 | 1 | 1 | 1 | 0.198337577 | 1 | 0.00049975 |
| 1 | 0.018997025 | 1 | 1 | 0.072140653 | 1 | 9.38E-08 | 1 | 0.00049975 |
| 1 | 0.004801254 | 1 | 1 | 1 | 1 | 1 | 1 | 0.0009995 |
| 0.094959214 | 1.12E-13 | 0.408311077 | 0.120712576 | 4.76E-11 | 0.236882141 | 6.05E-34 | 0.184700654 | 0.00049975 |
| 0.614440733 | 7.89E-06 | 1 | 1 | 1 | 1 | 1 | 1 | 0.00049975 |
| 0.550139302 | 0.055262777 | 0.013961441 | 0.226881856 | 1 | 1 | 0.000326551 | 0.234878832 | 0.00049975 |
| 1 | 1 | 1 | 1 | 1 | 1 | 1 | 1 | 0.00049975 |
| 0.094959214 | 1 | 1.61E-08 | 0.080087094 | 5.32E-20 | 1 | 1.18E-10 | 0.101536753 | 0.00049975 |
| 1 | 0.040645314 | 1 | 1 | 1 | 1 | 1 | 1 | 0.00149925 |
| 1 | 1 | 1 | 1 | 1 | 1 | 1 | 1 | 0.00049975 |
| 1 | 1 | 0.019868964 | 1 | 1 | 1 | 5.61E-05 | 1 | 0.00049975 |
| 1 | 1 | 1 | 1 | 1 | 1 | 1 | 1 | 0.00049975 |
| 1 | 0.1038879 | 1 | 1 | 1 | 1 | 1.20E-05 | 1 | 0.00049975 |
| 8.00E-142 | 0 | 1 | 1 | 1 | 0.004332651 | 8.19E-92 | 0.018447944 | 0.00049975 |

**Supplementary file**

COVID-19: Variant screening, an important step towards precision epidemiology

Amrita Chattopadhyay[1], Tzu-Pin Lu[1,2], Ching-Yu Shih[1], Liang-Chuan Lai[1,3], Mong-Hsun Tsai[1,4,5], Eric Y. Chuang[1,6,7*]

[1]Bioinformatics and Biostatistics Core, Centre of Genomic and Precision Medicine, National Taiwan University, Taipei 10055, Taiwan

[2]Institute of Epidemiology and Preventive Medicine, National Taiwan University, Taipei 10055, Taiwan

[3]Graduate Institute of Physiology, National Taiwan University, Taipei 10051, Taiwan

[4]Institute of Biotechnology, National Taiwan University, Taipei 10672, Taiwan

[5]Center of Biotechnology, National Taiwan University, Taipei 10672, Taiwan.

[6]Graduate Institute of Biomedical Electronics and Bioinformatics, National Taiwan University, Taipei 10617, Taiwan

[7]Biomedical Technology and Device Research Laboratories, Industrial Technology Research Institute, Hsinchu, Taiwan

*To whom all correspondences should be addressed

Eric Y. Chuang

Department of Electrical Engineering, Graduate Institute of Biomedical Electronics and Bioinformatics, National Taiwan University, Taipei 10617, Taiwan, Phone: +886-2-3366-3660, Fax: +886-2-3366-3682,

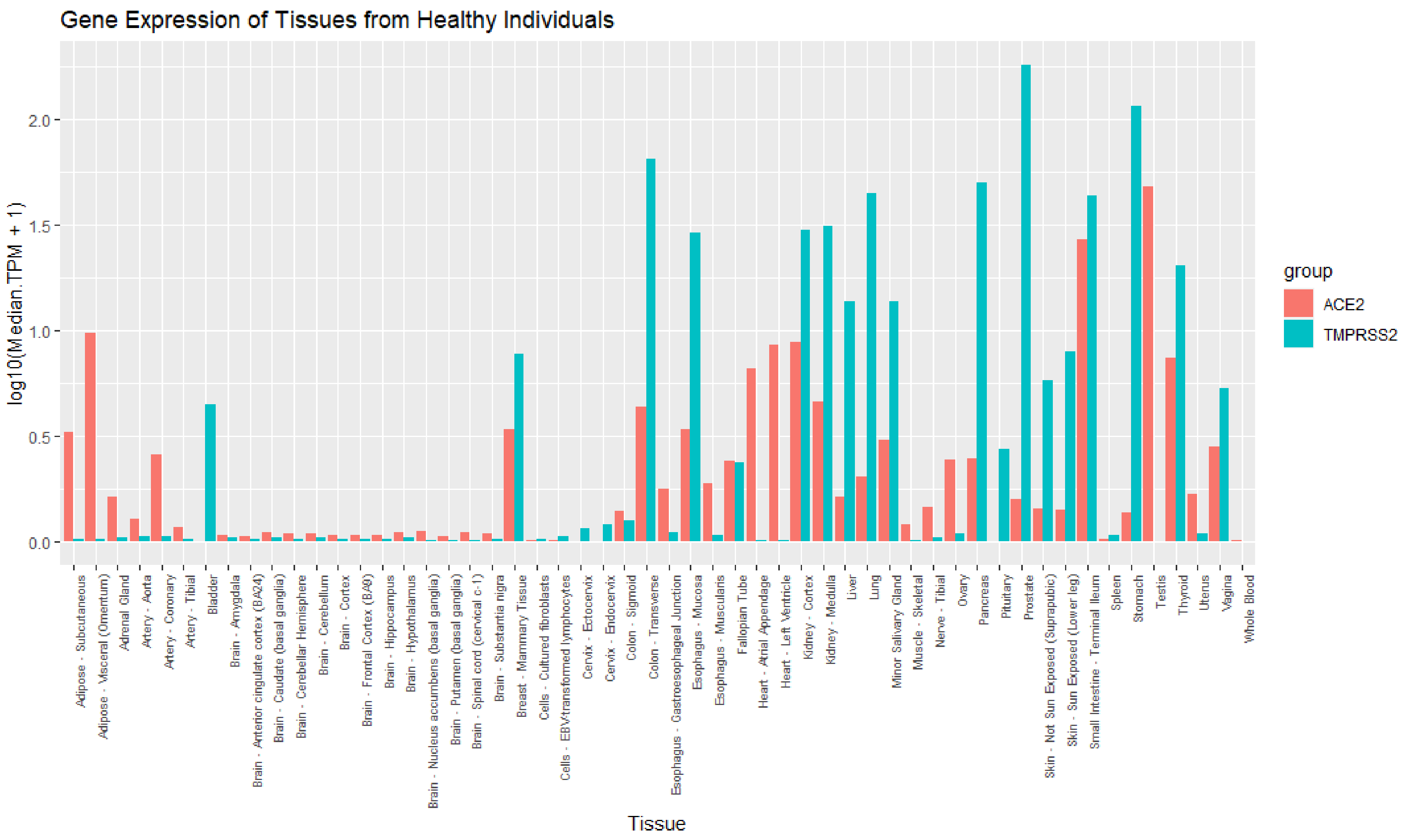


Supplementary file 3. Tissue specific gene expression in the GTEx healthy population

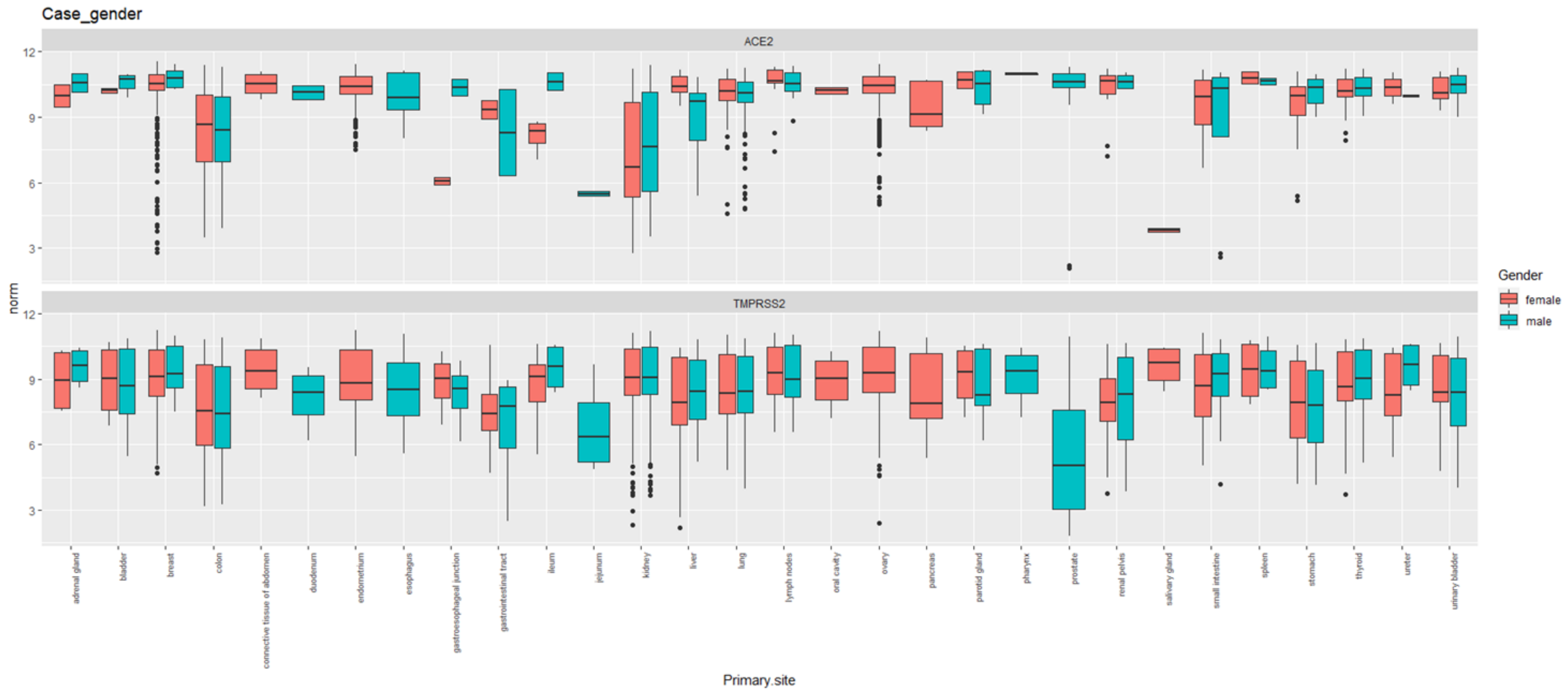


**Supplementary File 4. Gene expression of patients with cancers in different tissues from CellExpress, stratified by gender.** The x-axis displays tissue names; the y-axis displays norm = $\log_2$ normalized gene expression values. Pink box plots display gene expression for females; blue box plots display gene expression for males. The upper panel shows gene expression for *ACE2*; the lower panel shows gene expression for *TMPRSS2*. All gene expression data were downloaded from CellExpress (http://cellexpress.cgm.ntu.edu.tw/).

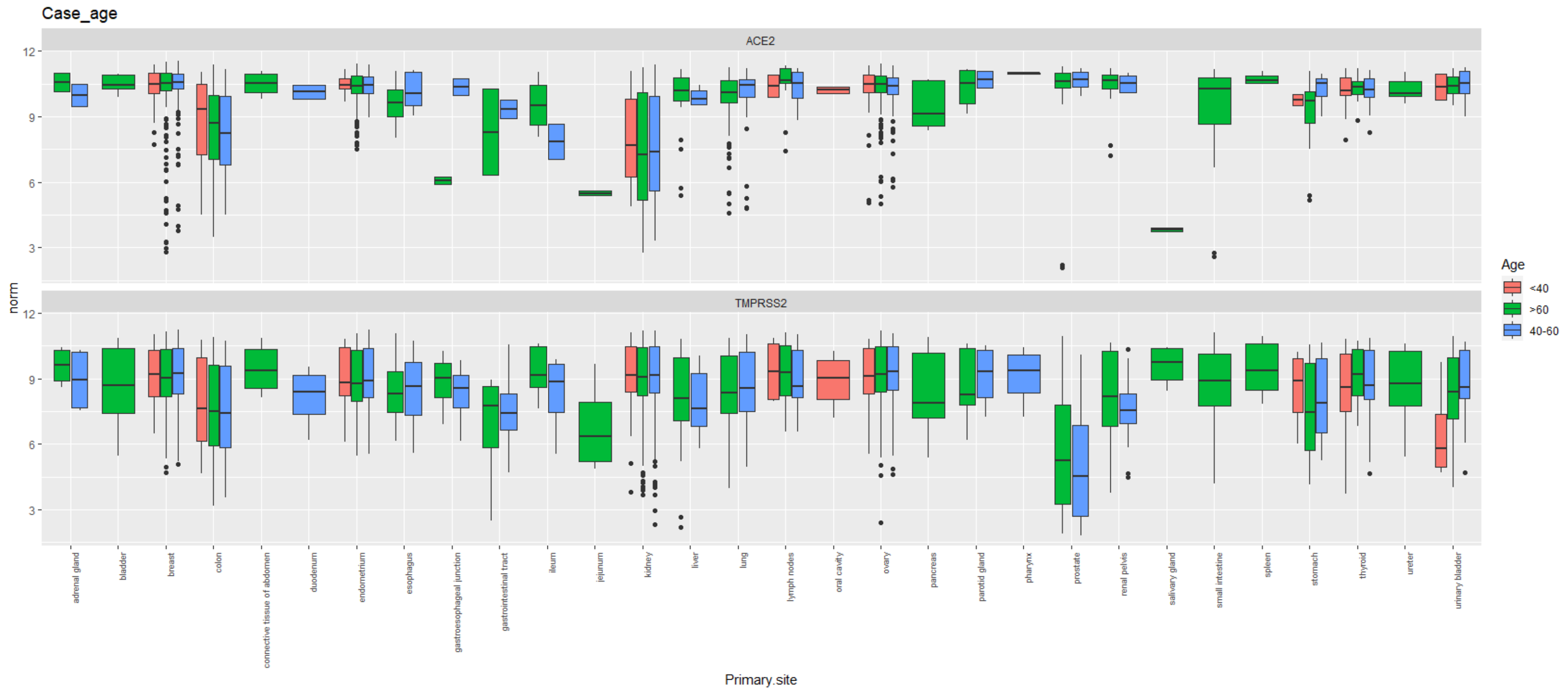


**Supplementary file 5. Gene expression of patients with cancers in different tissues from CellExpress, stratified by age.** The x-axis displays tissue names; the y-axis displays norm = $\log_2$ normalized gene expression values. Pink box-plots display gene expression for people with age <40 years; blue box-plots display gene expression for people with age >40 years and <60 years; and green box-plots display gene expression for people with age >60 years. The upper panel shows gene expression for *ACE2*; the lower panel shows gene expression for *TMPRSS2*. All gene expression data were downloaded from CellExpress (http://cellexpress.cgm.ntu.edu.tw/).
